# BRG1 generates subnucleosomes that expand OCT4 binding and function beyond DNA motifs at enhancers

**DOI:** 10.1101/2022.09.15.507958

**Authors:** Marina C. Nocente, Anida Mesihovic Karamitsos, Emilie Drouineau, Waad Albawardi, Cécile Dulary, Florence Ribierre, Hélène Picaud, Olivier Alibert, Joël Acker, Jean-Christophe Aude, Nick Gilbert, Françoise Ochsenbein, Sophie Chantalat, Matthieu Gérard

## Abstract

BRG1, the catalytic subunit of the mammalian SWI/SNF complexes, is essential for chromatin opening at enhancers. However, the nature of the open chromatin remains unclear. Here we show that in addition to producing histone-free DNA, BRG1 generates hemisome-like subnucleosomal particles containing the four core histones associated with 50-80 base pairs of DNA. Our genome-wide analysis indicates that BRG1 makes these particles by targeting and splitting fragile nucleosomes. In mouse embryonic stem cells, these subnucleosomes become an *in vivo* binding substrate for the master transcription factor OCT4 independently of the presence of OCT4 DNA motifs. At enhancers, the OCT4-subnucleosome interaction increases OCT4 occupancy and amplifies the genomic interval bound by OCT4 by up to one order of magnitude, compared to the region occupied on histone-free DNA. We suggest that BRG1-dependent subnucleosomes orchestrate an epigenetic mechanism that projects OCT4 function in chromatin opening beyond its DNA motifs.

## Introduction

High throughput analysis of chromatin organization and transcription factor (TF) distribution across the genome has led to considerable advances in our knowledge of the mechanisms regulating gene expression in mammalian cells. Despite these advances, understanding how transcription factors (TFs) recognize their target cis-regulatory elements (CREs) on the mammalian genome and regulate transcription remains a major challenge^1,2^. Most mammalian TFs bind transiently to short (5-15 bp) and often degenerated DNA motifs^3,4^. Due to its large size, the genome provides non-functional binding opportunities that largely exceed the number of interaction events experimentally detected at CREs^5,6^. The nucleosomal organization and its dynamic regulation are the main mechanisms that orientate the binding of TFs to their proper targets on the genome.

The packaging of the genome into nucleosomes is a physical barrier to the binding of TFs and enzymes^2,7^. Nucleosomes are made of a core composed of two copies of each of the histones H2A, H2B, H3 and H4, around which 147 bp of DNA form two gyres interacting with the surface of the core^8^. The rotational positioning of the motifs on the DNA helix impairs TF binding when the motif faces the core^9^. In addition, the motifs of most TFs are too long to fit entirely on the solvent-exposed DNA. This last constraint can be overcome by TFs with pioneer activity^9^ such as OCT4, a master TF that controls pluripotency and self-renewal of mouse embryonic stem cells (mESCs)^10^. OCT4 can interact with partial motifs on the solvent-exposed side of the DNA, showing that nucleosomes can also support specific TF-DNA interactions on the genome^11,12^. However, OCT4 interacts more efficiently with partial motifs located near the DNA entry/exit site, where nucleosomal DNA is more loosely bound^13,14^, showing that the obstructive role of the nucleosome balances this supportive function.

The nucleosomal barrier is dynamically regulated by a large family of enzymes called chromatin remodelling factors (remodelers), which modulate nucleosome positioning and occupancy at CREs^15–18^. In particular, the SWI/SNF (switch/sucrose non-fermentable) complexes (also called BAF complexes), which bear tumor suppressor properties^19^, are essential for generating accessible DNA regions at enhancers^20–22^. It is generally assumed that the main product of SWI/SNF-mediated chromatin remodeling corresponds to DNA stripped of histones and that this histone-free DNA is the main genomic template bound by TFs at CREs *in vivo*. However, some TFs can also interact with unstable, micrococcal nuclease (MNase)-sensitive nucleosomes (also called fragile nucleosomes) at CREs^23,24^. In yeast, these fragile nucleosomes correspond to partially unwrapped nucleosomes generated by the RSC (remodeling the structure of chromatin) complex^24^, which is an ortholog of the mammalian SWI/SNF complexes.

In this study, we investigated the function of BRG1 (SMARCA4), the ATPase of SWI/SNF complexes in mESCs, using a method that distinguishes histone-free DNA from histone-containing genomic particles among the products of chromatin remodeling. As well as generating histone-free DNA, we show that BRG1 produces hemisome-like subnucleosomal particles encompassing 50-80 bp of DNA at enhancer elements. We further demonstrate that these subnucleosomal particles are an *in vivo* binding substrate for OCT4 in mESCs, independently of the presence of OCT4 motifs in the DNA. This interaction with the 50-80 bp subnucleosomes increases OCT4 occupancy and allows a striking expansion of the genomic interval occupied by OCT4 relative to the short interval it occupies on histone-free DNA. We also show that OCT4’s potent function in chromatin opening at enhancers^20^ is active precisely within the genomic interval delimited by the interaction between OCT4 and the 50-80 bp subnucleosomes. Together, these results reveal an epigenetic mechanism based on BRG1-generated subnucleosomal particles that expand OCT4’s binding interval and project OCT4’s function beyond the boundaries of its DNA motifs.

## Results

### Enhancers have a discrete nucleosomal and subnucleosomal organization

In mESCs, enhancers are bound by a series of TFs that are required to maintain pluripotency and self-renewal, including OCT4, SOX2, NANOG, ESRRB and KLF4^25^. The DNA binding sites (BS) for these TFs are located in relative proximity to each other, yet without a constrained arrangement pattern^3,25–28^. Enhancers are also characterized by their hypersensitivity to DNase I, the presence of the mediator complex, and, for active enhancers, RNA polymerase II (Pol II) and TATA box Binding Protein (TBP), as well as H3K27ac^29–32^. The chromatin organization of enhancers is less well defined than that of promoters and CCCTC-binding factor (CTCF)-bound loci^23,33–39^. We investigated nucleosomal organization at enhancers by micrococcal nuclease digestion of chromatin, followed by immunoprecipitation using antibodies against histone H3 and deep sequencing (MNase ChIP-seq) (**Fig. 1a**). We show that k-means clustering based on the distribution patterns of canonical nucleosomes and subnucleosomal-size particles leads to identifying clusters of enhancers with different but highly related chromatin organizations (Extended Data Fig. 1a).

**Figure 1.**
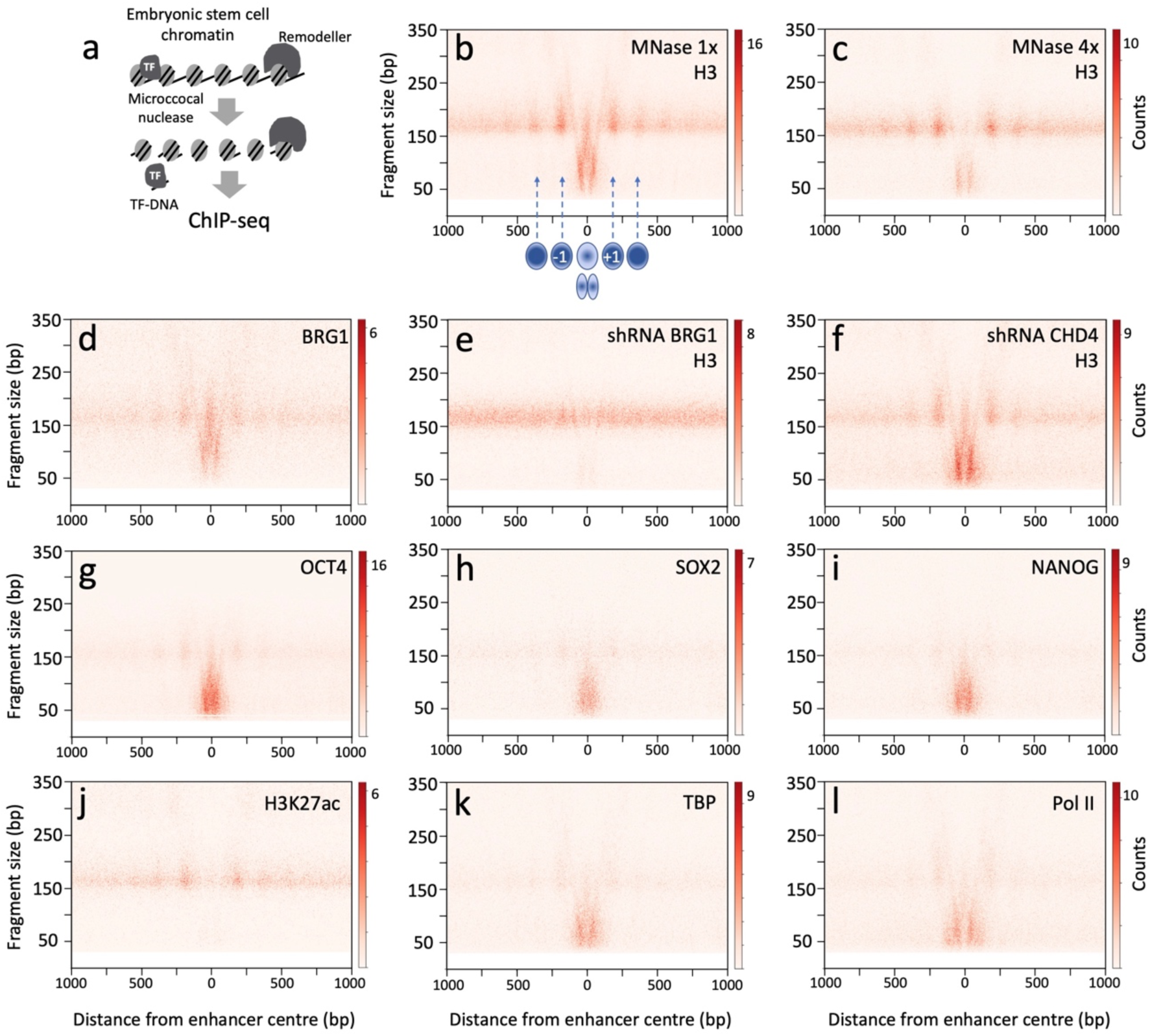
BRG1 controls nucleosomal and subnucleosomal organization at enhancers. **a**, MNase-digested chromatin was prepared from mESCs for ChIP-seq experiments. **b**, **c**, V-plots of histone H3 ChIP-seq fragments spanning ± 1000 bp from cluster 1 enhancer center, using either the standard MNase dose (**b**) or a four-fold excess (**c**). Red dots indicate on the *x-axis* the genomic position of the midpoint of each immunoprecipitated DNA fragment and its length in bp on the *y-axis*. The color scale corresponds to the number of DNA fragments. The schematic illustration in (**b**) indicates the positions of nucleosomes −1 and +1, which flank the central fragile nucleosome and associated subnucleosomal particles. **d,** V-plot of BRG1 ChIP-seq. **e**, **f**, V-plots of histone H3 ChIP-seq in mESCs depleted of either BRG1 (**e**) or CHD4 (**f**), using shRNAs. **g**-**l**, V-plots of OCT4, SOX2, NANOG, H3K27ac, TBP and Pol II ChIP-seq. The standard MNase dose was used in all panels except in (**c**).

Using two-dimensional maps (V-plot)^40^, we detected in the central region of each cluster of enhancers the presence of short (50-120 bp) DNA fragments associated with histone H3, corresponding to subnucleosomal particles precisely positioned across the enhancers (Extended Data Fig. 1b). These subnucleosomal particles alternate in position with one or more full-size nucleosomes (i.e., histone-associated 150-180 bp long DNA fragments), depending on the distance that separates the two well-positioned nucleosomes, defined hereafter as nucleosomes −1 and +1, that flank the center of the enhancer (**Fig. 1b** and Extended Data Figs. 1b, 2a). Enhancer clusters 1 to 3 that are representative of all subgroups differ from each other according to the distance between nucleosomes −1 and +1, with cluster 1 having the narrowest (340 bp) and cluster 3 the widest (> 480 bp) distance (Extended Data Fig. 1b).

In enhancer cluster 1 and related clusters 5, 8 and 12, a single centrally located nucleosome is present between −1 and +1 nucleosomes (**Fig. 1b** and Extended Data Fig. 1b). This central nucleosome is highly sensitive to MNase dosage, as it completely disappeared when mESCs were incubated with a four-fold excess of MNase (**Fig. 1c**), and we will hereafter refer to this particle as a fragile nucleosome^23^. In contrast, the flanking nucleosomes −1 and +1 were only slightly affected by the high-dose MNase treatment, revealing their canonical nature. Enhancer cluster 2, and related clusters 4, 9 and 10, are characterized by the presence of fragile nucleosomes at two main positions instead of one in the central interval between nucleosomes −1 and +1 (Extended Data Fig. 2a, b). We observed that in cluster 1, the single central fragile nucleosome is symmetrically flanked on each side of its dyad axis by a subnucleosomal particle protecting from MNase digestion a minimum DNA length of 54 bp, as assessed by the length of the DNA fragments present at the vertex of the ‘V’ shape signal (**Fig. 1b** and Extended Data Fig. 3a). In cluster 2, each of the two fragile nucleosomes of the central interval is flanked by subnucleosomal particles in a pattern similar to cluster 1 (Extended Data Fig. 2a and Extended Data Fig. 3a). Enhancer cluster 3 shows a variation of these patterns, with a wider central interval accommodating a larger number of subnucleosomal particles (Extended Data Fig. 1b). Thus, our analysis revealed that enhancers have a modular chromatin organization in which two positioned canonical nucleosomes delimit a central region containing a variable number of basic modules. Each basic module comprises one fragile nucleosome flanked on both sides of its dyad axis by two subnucleosomal particles occupying the same genomic interval, suggesting alternating chromatin configurations.

### Enhancer subnucleosomal particles are enriched in TF binding motifs

To investigate the relation between this discrete chromatin architecture and TF motif distribution, we mapped the DNA-interacting motifs of the pluripotency-associated TFs OCT4, SOX2, NANOG, and KLF4^25^. The genomic interval encompassing the central fragile nucleosome(s) and associated-subnucleosomal particles corresponds precisely to the area in which TF motifs are enriched (Extended Data Fig. 3b, c). Thus, the larger size of this genomic interval at multi-modular enhancers correlates with a greater number of OCT4 and other TF motifs compared to single-module enhancers (Extended Data Fig. 3). We then examined the distribution of TFs and epigenetic features that contribute to enhancer functions (**Fig. 1g-l** and Extended Data Fig. 2f-k). ChIP-seq analysis revealed that all tested TFs bind within the central interval of the enhancer in which subnucleosomal particles are interspersed with fragile nucleosome(s). Even TFs known to have pioneer functions, such as OCT4 and SOX2, were preferentially enriched on DNA fragments spanning 50 to 120 bp in length rather than the 150-180 bp typical of canonical nucleosomes (**Fig. 1g, h** and Extended Data Fig. 2f, g). The general transcription factor TBP and Pol II were associated with 40-120 bp DNA fragments within the same central interval of the enhancer occupied by OCT4 and SOX2 (**Figure 1k, l** and Extended Data Fig. 2i, k). In contrast to nucleosomes −1 and +1, the subnucleosomal particles were unexpectedly devoid of H3K27ac signal (**Fig. 1j** and Extended Data Fig. 2j). The enhancers selected for this study lack CTCF ChIP-seq signal (Methods), excluding the possibility that this TF might organize nucleosome positioning.

### BRG1 controls nucleosomal and subnucleosomal organization at enhancers

We then investigated which factors control this enhancer-specific nucleosomal organization. ATP-dependent chromatin remodelers^15–18^ are prime candidates with BRG1 in particular forming accessible chromatin at enhancers^20–22^. Depletion of BRG1 using an shRNA resulted in striking alterations of both nucleosomal and subnucleosomal organization at enhancers. Subnucleosomal particles were almost completely absent at enhancers in cells depleted of BRG1, compared to control cells (**Fig. 1e** and Extended Data Fig. 2d). In addition, the positioning of canonical nucleosomes was severely altered, resulting in their apparently random redistribution across the enhancer. We detected identical alterations with antibodies against histone H3 and H2B, as well as when we used a second shRNA targeting a different region of BRG1 mRNA (Extended Data Fig. 4a). These perturbations of chromatin organization at enhancers were specific to BRG1 loss-of-function, as shRNA-mediated depletion of chromatin remodeler CHD4 did not prevent the generation of subnucleosomal particles, nor the positioning of canonical nucleosomes (**Fig. 1f** and Extended Data Fig. 2e). Depletion of BRG1 using the auxin-induced degron (AID) system^41^ resulted in similar defects in chromatin organization even after a short period of auxin treatment, showing that impairment of subnucleosomal organization is not a secondary effect of BRG1 loss-of-function (Extended Data Fig. 4b, c).

The alterations in chromatin organization caused by BRG1 depletion are compatible with a scenario in which BRG1 binds the central fragile nucleosome(s) to convert it into smaller particles, such as hexasomes [(H3-H4)_2_; H2A-H2B] or hemisomes (half-nucleosomes) containing one copy of each of the histones H3, H4, H2A, H3B^42–45^. In support of this hypothesis, we detected BRG1 enriched both at the centrally located fragile nucleosome(s) and at subnucleosomal particles (**Fig. 1d** and Extended Data Fig. 2c). In an alternative hypothesis, BRG1 could extrude a DNA loop on the central nucleosome and propagate it to the dyad axis region^46^, which would become highly sensitive to MNase digestion and yield a histone octasome associated with two short DNA fragments symmetrically positioned on each side of the dyad, instead of the regular 150-180 bp-long fragment. To test these hypotheses, we set up an experimental strategy based on separating nucleosomal and subnucleosomal particles by centrifugation through a sucrose gradient, in which the particles sediment as a function of their molecular mass and shape^47^ (**Fig. 2a**). We loaded the MNase-digested mESC chromatin onto a 10-30% sucrose gradient. After centrifugation, individual fractions were collected and analyzed by agarose gel electrophoresis, revealing genomic DNA fragments ranging from 30 to 120 bp, which migrated slower than canonical nucleosomes through the gradient (**Fig. 2b**). To identify which fractions contained subnucleosomal particles, we performed ChIP-seq experiments with an antibody against histone H3 (**Fig. 2c**). At the top of the gradient, the DNA fragments in fractions 3-4 were not efficiently immunoprecipitated, suggesting that they mostly correspond to histone-free DNA fragments. In contrast, the 50-80 bp-long particles collected in gradient fractions 5-6, which have a low sedimentation rate, were associated with histone H3 at enhancers and are thus genuine subnucleosomal particles (**Fig. 2c**). Their slow sedimentation rate is incompatible with the hypothesis that these particles correspond to histone octamers associated with two 50-80 bp DNA fragments. Such particles would have approximately the same molecular weight as canonical nucleosomes and thus would be expected to sediment in fractions 11 to 13.

**Figure 2.**
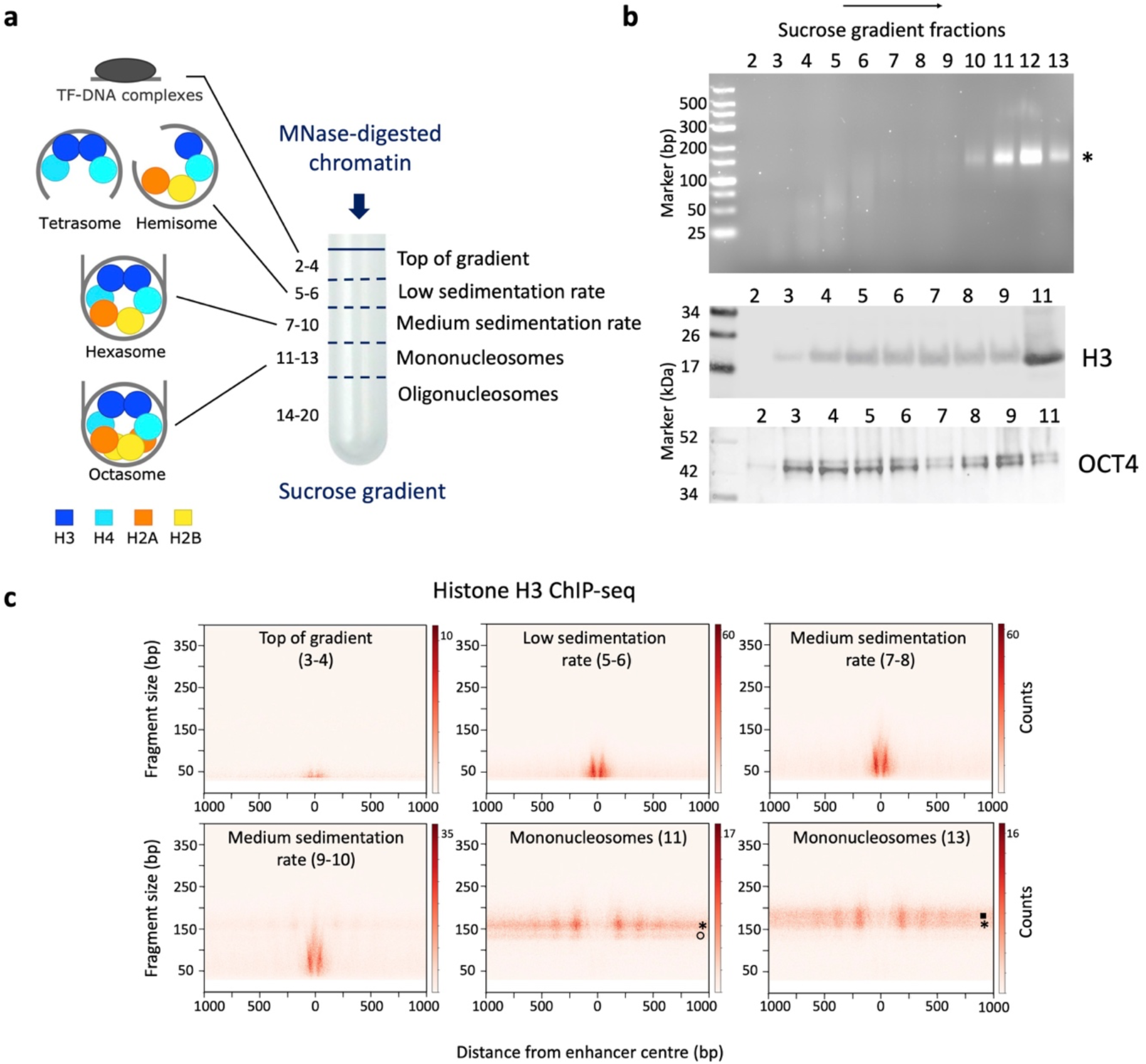
Identification of low sedimentation rate subnucleosomal particles associated with 50-80 bp of DNA at enhancers. **a**, MNase-digested chromatin was centrifugated through a 10-30% sucrose gradient to separate canonical nucleosomes from putative subnucleosomal particles and TF-DNA complexes. **b**, The DNA fragments of each fraction were analyzed by electrophoresis in a 4% agarose gel (top), and the distribution of histone H3 and OCT4 was revealed by Western blot (mid and bottom). The asterisk indicates the DNA fragments corresponding to mononucleosomes. **c**, V-plots of histone H3 ChIP-seq experiments performed with sucrose gradient fractions as input, spanning ± 1000 bp from cluster 1 enhancer center. The fractions or pools used for ChIP are indicated on each panel. The open circle and black square point to potential partially unwrapped nucleosomes and chromatosomes, respectively.

### A subclass of subnucleosomal particles contains the four core histones associated with 50-80 bp of DNA

To define the histone content of the low sedimentation rate 50-80 bp subnucleosomal particles (hereafter called 50-80 bp subnucleosomes), we performed ChIP-seq experiments with antibodies against the remaining three histones H4, H2A and H2B. These antibodies efficiently immunoprecipitated the 50-80 bp subnucleosomes but not the DNA fragments from the top of the gradient (**Fig. 3**). This result excludes the possibility that the 50-80 bp subnucleosomes might correspond to tetrasomes composed of two dimers of histones H3 and H4, which are nucleosome assembly intermediates detected during DNA replication^48^. The presence of the four core histones suggests that these particles correspond to either hexasomes or hemisomes^42,44,45,49^. However, the DNA length (50-80 bp) protected from MNase digestion is incompatible with the ~90 bp DNA associated with hexasomes^50^. The protection from MNase cleavage conferred by hexasomes yielded DNA fragments ranging from 90 to 130 bp^42^. In our experimental setting, subnucleosomal particles associated with 90-130 bp DNA fragments have a medium sedimentation rate (fractions 7-8 and 9-10), suggesting a histone composition different from the 50-80 bp subnucleosomes that sedimented in fractions 5-6 (**Fig. 2**). Furthermore, the positions of the DNA fragment centers of these 50-80 bp particles are symmetrically distributed on each side of the dyad axis of the central fragile nucleosome, and the distance spanning the sum of the lengths of the two consecutive particles (140-180 bp) is equivalent to the size of DNA wrapped around an octasome (Extended Data Fig. 3a). A comprehensive examination of individual single-module enhancers from clusters 1 and 8 revealed that 50-80 bp subnucleosomes are systematically associated in pairs and separated by a DNA linker having a median length of 29 bp (**Fig. 4d**, Extended Data Fig. 5 and Methods). This unique genomic positioning of the 50-80 bp subnucleosomes, their histone content, and their size properties suggest that they correspond to split nucleosomes that are composed of two hemisomes connected by a linker DNA fragment of variable length. This linker DNA is likely hypersensitive to MNase digestion, which separates the two hemisomes from each other and results in their co-segregation within the low sedimentation rate fractions of the gradient. We verified the systematic co-segregation of the four histones H2A, H2B, H3 and H4 in 50-80 bp subnucleosomes by observing a large number (n = 2,686) of individual enhancers (Extended Data Fig. 5 and Methods). We confirmed that the 50-80 subnucleosomes have a molecular size markedly inferior to mononucleosomes using size exclusion chromatography (Extended Data Fig. 6a-c).

**Figure 3.**
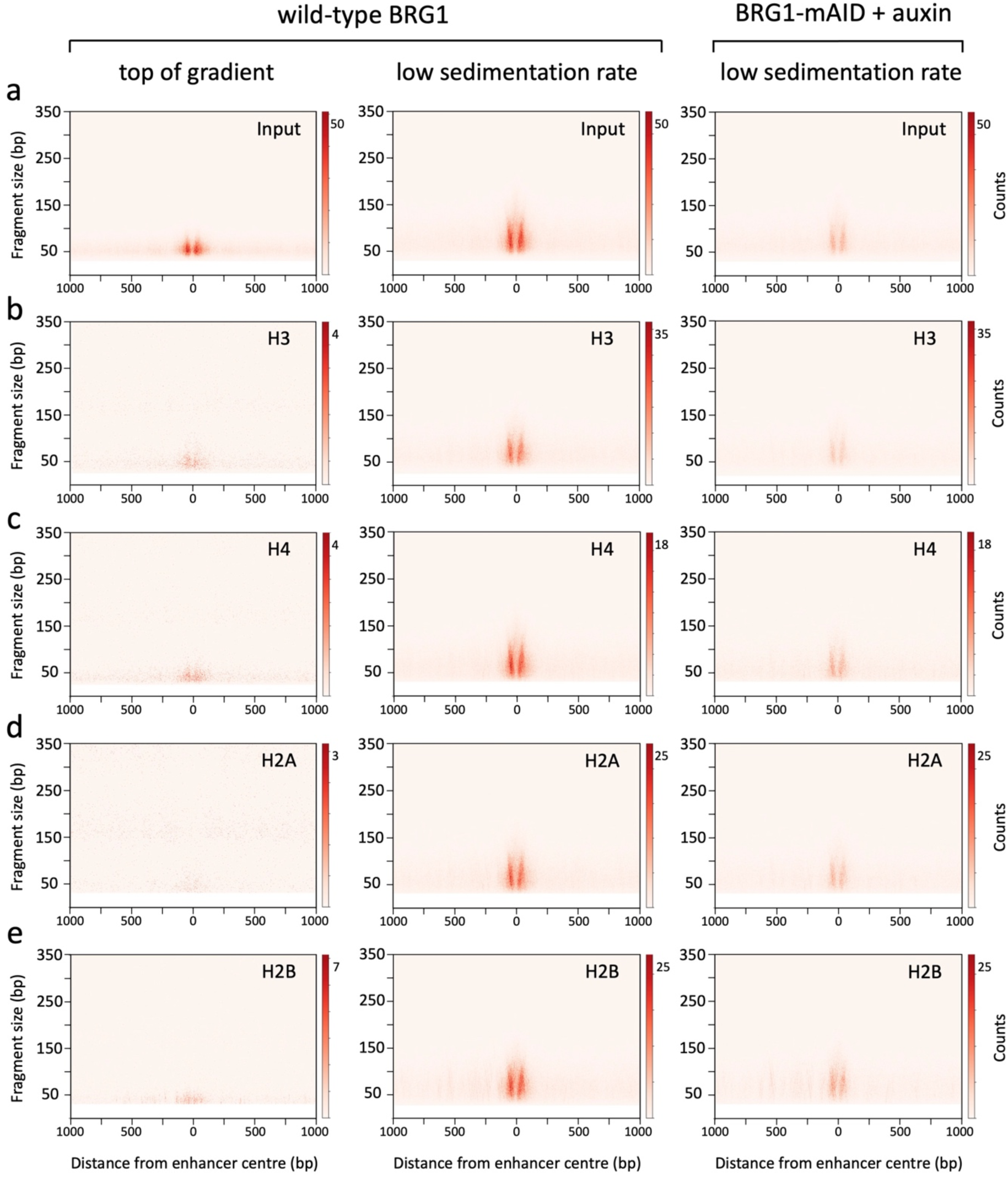
Histone composition of 50-80 bp subnucleosomes. Chromatin, prepared from mESCs expressing wild-type BRG1, or from cells depleted of BRG1 using the AID system, was centrifugated through a sucrose gradient as in Fig. 2. **a**, V-plots of the DNA fragments which sedimented at the top of the gradient (pool of fractions 2-3-4), or of the DNA fragments present in low sedimentation rate fractions (pool of fractions 5-6), spanning ± 1000 bp from the cluster 1 enhancer center. Signals shown in the two right panels were adjusted according to the total number of fragments sequenced in each dataset. **b-e**, V-plots of histone H3, H4, H2A and H2B ChIP-seq experiments performed with the pools of sucrose gradient fractions indicated in (**a**) as input. For each histone, the signals of the ChIP-seq experiments performed with subnucleosomal particles from mESCs expressing wild-type BRG1 and from BRG1-depleted cells were adjusted according to the total number of DNA fragments in each dataset.

**Figure 4.**
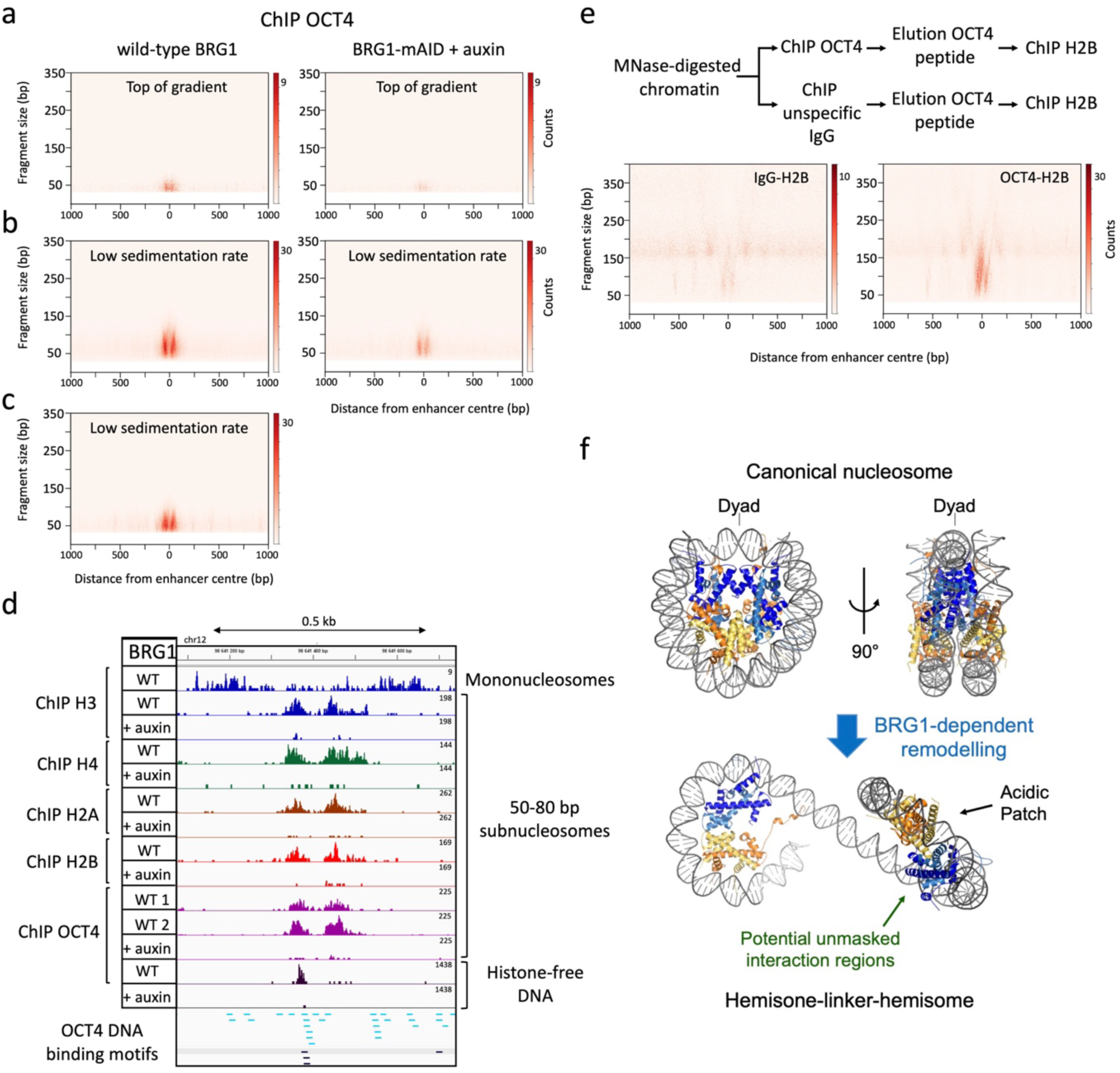
OCT4 binds to 50-80 bp subnucleosomes at enhancers. **a**-**c**, V-plots of ChIP-seq fragments spanning ± 1000 bp from the cluster 1 enhancer center. Chromatin, prepared from mESCs expressing wild-type BRG1 (left panels) or from cells depleted of BRG1 using the AID system (right panels), was centrifugated through a sucrose gradient as in Fig. 2. OCT4 ChIP experiments were performed using DNA and chromatin fragments which sedimented at the top of the gradient (**a**, pool of fractions 2-3-4), or present in low sedimentation rate fractions (**b**, **c**, pool of fractions 5-6). Two distinct OCT4 antibodies were used in (**b**) and (**c**). **d**, Density graphs showing the distribution of ChIP DNA fragment centers at a representative example of a cluster 1 enhancer. The top lane shows the positions of canonical nucleosomes detected by histone H3 ChIP-seq of sucrose gradient fractions 11 and 13. The following eight lanes show the distribution of 50-80 bp subnucleosomes, revealed by histone H3, H4, H2A and H2B ChIP-seq of sucrose gradient fractions 5-6. The following five lanes show the OCT4 ChIP-seq signal on 50-80 bp subnucleosomes or histone-free DNA from the top gradient fractions. Experiments were performed with chromatin of cells expressing wild-type (WT) BRG1 or depleted of BRG1 using the AID system (+ auxin). Two distinct antibodies against OCT4 were used in lanes WT 1 and 2. High (p = 0.001) and low (p = 0.01) confidence OCT4 consensus binding motifs are indicated in black and blue, respectively. **e**, Sequential ChIP-seq experiments were performed on chromatin prepared from mESCs expressing wild-type BRG1. Chromatin was first immunoprecipitated with an OCT4 antibody or with an unspecific IgG. Elution was performed by peptide competition, and each eluted fraction was subjected to a second round of immunoprecipitation with an antibody against H2B. The right and left panels show the sequential OCT4-H2B ChIP-seq and the unspecific IgG control experiment. **f**, Model proposing that BRG1 splits nucleosomes at enhancers to generate hemisome-linker-hemisome entities, which might display new interaction regions for TFs. The two hemisomes are separated by a DNA linker having a median length of 29 bp. Histones H3 and H4 are shown in two shades of blue, and histones H2A and H2B are in yellow and orange, respectively.

We next used transmission electron microscopy (EM) to analyze the size of 50-80 bp subnucleosomes compared to mononucleosomes purified from sucrose gradients and nucleosomes assembled *in vitro* onto an array of 601 DNA fragments^51^ (Extended Data Fig. 6d, e). Canonical nucleosomes have a front view diameter of 10 to 13 nm and a width of about 6 nm, the latter of which is defined by the sum of the two DNA gyres wrapped around the histone octamer^52^. If the particles generated at enhancers are hemisomes, they should retain a 10-13 nm diameter but have a width reduced to 3 nm, which corresponds to the size of a single DNA gyre. EM analysis should then reveal particles having an apparent size either smaller or equal to that of canonical nucleosomes, depending on their orientation on the grid. Random selection and size measurement of both particles from gradient fractions 5-6 revealed a large proportion of small-sized particles (< 10 nm), which are absent in control nucleosome-containing fractions (Extended Data Fig. 6d, e). The EM analysis thus further supports the hypothesis that enhancer 50-80 bp subnucleosomes correspond to hemisomes.

We then considered whether the 50-80 bp subnucleosomes are specific to mESCs or if they are also produced in other mammalian cells. We used the histone MNase ChIP-seq approach combined with sucrose gradient sedimentation experiments to probe chromatin organization in a human melanoma cell line. We isolated clusters of putative enhancers enriched for 50-80 bp subnucleosomes, which have a chromatin organization very similar to that of mESCs enhancer clusters (Extended Data Fig. 7). This analysis shows that the modular organization of nucleosomes and subnucleosomes we described at mESCs enhancers is conserved in human cells.

### 50-80 bp subnucleosomal particles are produced at all categories of CREs

We used our high-coverage histone MNase ChIP-seq datasets to explore subnucleosomal organization at CTCF-BS and promoters. This analysis revealed the presence of 50-80 bp subnucleosomes having distribution patterns distinct from that of enhancers (Extended Data Fig. 8). At CTCF-BS, a single 50-80 bp subnucleosome was detected in the central region coinciding with the location of the CTCF motif (Extended Data Fig. 8a, b). Promoters contain a single 50-80 bp subnucleosome coinciding in genomic position with the nucleosome −1, located upstream of the TSS (Extended Data Fig. 8c, d). This nucleosome −1 was hypersensitive to an excess of MNase (Extended Data Fig. 8c), a hallmark of fragile nucleosomes. These observations suggest that at promoters, chromatin remodeling events target nucleosome −1 to convert it into a 50-80 bp subnucleosome occupying the same genomic position. In support of this hypothesis, we have shown previously that a large variety of chromatin remodelers bind nucleosome −1 in mESCs^18^. We detected a 30 bp protection against MNase digestion (footprint) within the 50-80 bp subnucleosome of promoters, which matches precisely the 30 bp of DNA upstream of the TSS. This footprint reveals the binding on the subnucleosome of proteins potentially involved in preinitiation complex (PIC) formation (Extended Data Fig. 8e). Since these particles are present in low sedimentation rate fractions, the footprint might be conferred by low molecular weight TFs like TBP and TFIIB but not by large complexes such as TFIID or Pol II.

### BRG1 generates the 50-80 bp subnucleosomal particles at enhancers

The chromatin perturbation pattern caused by BRG1 depletion (**Fig. 1**) suggests that this remodeler might generate the 50-80 bp subnucleosomes at enhancers. We tested this possibility by repeating the sucrose gradient sedimentation experiments, using chromatin prepared from either BRG1-depleted or control cells. ChIP-seq experiments with antibodies against histones H3, H4, H2A and H2B revealed that BRG1 depletion results in a strong reduction of the amount of 50-80 bp subnucleosomes, thus indicating that their production at enhancers is indeed dependent on BRG1 (**Fig. 3**). BRG1 depletion also reduced the amount of medium sedimentation rate 90-130 bp subnucleosomal particles, showing that BRG1 activity is required for the generation of all classes of subnucleosomal particles detected at enhancers in this study (Extended Data Fig. 9c-f). In contrast, BRG1 depletion did not alter the distribution or amount of subnucleosomes at CTCF-BS (Extended Data Fig. 8b). At promoters, BRG1 loss of function did not affect nucleosome positioning, but the amount of 50-80 bp subnucleosomes was slightly decreased (Extended Data Fig. 8c, d). Altogether, these results show that BRG1 activity is mainly required at enhancers to produce subnucleosomal particles and properly position nucleosomes.

We also tested whether BRG1 is involved in producing histone-free DNA fragments at enhancers. Deep sequencing of the DNA prepared from the top sucrose gradient fractions revealed that BRG1-depletion resulted in a pronounced reduction of these DNA fragments at enhancers but not at CTCF-BS (Extended Data Fig. 9a, b). DNA fragments containing motifs for OCT4, SOX2, ESRRB, KLF4 and NANOG were markedly depleted in the top gradient fractions prepared from BRG1-depleted cells (Extended Data Fig. 9g), demonstrating that BRG1 is required for the efficient production of histone-free, accessible DNA for these TFs at enhancers. BRG1 is thus essential at enhancers for producing two main chromatin remodeling products: histone-free DNA and 50-80 bp subnucleosomal particles.

A subset of enhancers is organized in clusters called super-enhancers, which contribute to the control of cell phenotype^53,54^. The nucleosomal and subnucleosomal organization of super-enhancers were identical to that of regular enhancers (Extended Data Fig. 10).

### 50-80 bp subnucleosomes interact with OCT4 independently of the presence of an OCT4 motif

We observed that in the OCT4, SOX2 and NANOG ChIP-seq experiments (**Fig. 1g-i**), the sizes and position on the genome of the immunoprecipitated DNA fragments were similar to that of 50-80 bp subnucleosomes (**Fig. 3**). This specific pattern, reproduced with three distinct antibodies against OCT4 (**Fig. 1, 4** and Methods), suggests that 50-80 bp subnucleosomes might represent a previously undetected binding substrate for TFs *in vivo*. To test if OCT4 is genuinely bound to histone-associated subnucleosomal particles, we carried out sequential ChIP experiments using antibodies against OCT4 in the first step. Elution of OCT4-bound chromatin was performed using a competitor peptide, and the eluted chromatin was used in a second step of ChIP with antibodies against histone H2B. Deep sequencing of the DNA associated with both OCT4 and histone H2B revealed DNA fragments of subnucleosomal size, thus demonstrating that OCT4 is bound to subnucleosomal particles *in vivo* (**Fig. 4e**). Analysis of OCT4-immunopurified 50-80 bp subnucleosomes by EM revealed particle sizes compatible with the hypothesis that OCT4 interacts *in vivo* with hemisomes (Extended Data Fig. 6d, e).

We next compared how OCT4 binds to 50-80 bp subnucleosomes and histone-free DNA present in the top gradient fractions. OCT4 antibodies successfully immunoprecipitated both histone-free DNA fragments and 50-80 bp subnucleosomes (**Fig. 4a, b**). However, OCT4 enrichment profiles were different at these two distinct genomic templates. At histone-free DNA fragments, OCT4 peaks coincided with OCT4 consensus motifs, whereas in 50-80 bp subnucleosomes, OCT4 enrichment coincided with histones (**Fig. 4d** and Extended Data Fig. 5, 10). The specificity of this unexpected OCT4 ChIP-seq enrichment profile on 50-80 bp subnucleosomes was verified with a second antibody against OCT4, and by performing the elution step of the ChIP experiment using a competitor peptide (**Fig. 4b-d**, and Extended Data Fig. 5, 10).

We observed this coincidence between OCT4 and histone enrichment even when OCT4 binding motifs were absent from the DNA wrapped around the particle (Extended Data Fig. 11 and Methods). It remained possible that partial OCT4 motifs could be enriched on subnucleosomes and compensate for the absence of full motifs, in a manner reminiscent of the interaction between OCT4 and the nucleosomes^11,12^. However, examination of the distribution of partial motifs did not reveal their enrichment on subnucleosomes, invalidating this hypothesis (Extended Data Fig. 11g and Methods). These observations suggest that OCT4 interactions with 50-80 bp subnucleosomes involve contacts with the histone component in addition to those with its DNA binding motif. OCT4 binding motifs are strongly enriched in the dyad region of the central fragile nucleosome of the enhancer (Extended Data Fig. 3b). BRG1-mediated conversion of this central nucleosome into two hemisome-like particles is associated with the appearance of a DNA linker between the two particles (Extended Data Fig. 5i). This linker contains the OCT4 motifs initially present in the dyad region of the central nucleosome. MNase treatment prior to ChIP-seq cuts away the linker; OCT4 would be released if it interacted with the subnucleosomal particle exclusively via its DNA motif. Since OCT4 remains bound to hemisome-like particles that do not contain an OCT4 motif (Extended Data Fig. 11), we concluded that these particles must provide an interface stabilizing OCT4 interaction. An intriguing possibility is that the internal histone surface of the split nucleosome, which is inaccessible in canonical nucleosomes, provides an interaction domain for OCT4 and potentially for other TFs and chromatin-binding proteins in a manner reminiscent of the acidic patch of the nucleosome, which is present on the other side (**Fig. 4f**).

### 50-80 bp subnucleosomes expand the genomic interval bound by OCT4 and increase its occupancy at enhancers

To address the function of the interaction between OCT4 and 50-80 bp subnucleosomes, we first compared the size of the genomic intervals bound by OCT4 at either 50-80 bp subnucleosomes or histone-free DNA. This comparison revealed an up to seven-fold size increase for 50-80 bp subnucleosomes relative to histone-free DNA (**Fig. 5a-d**). Thus, the function of the interaction between OCT4 and 50-80 bp subnucleosomes is to dramatically increase the size of the genomic interval bound by OCT4 at enhancers. A second potential function of the 50-80 bp subnucleosomes could be to augment OCT4 occupancy of enhancers. A previous single-molecule footprinting analysis revealed a low OCT4 occupancy on nucleosome-free DNA molecules bearing its consensus motif in mESCs^3^. In agreement with this result, we observed that about 10% of the enhancers display a robust OCT4 ChIP-seq signal on histone-free DNA (**Fig. 4d**, **5i**, left panel, Extended Data Fig. 5e), whereas 90% have a low or undetectable signal (**Fig. 5j**, Extended Data Fig. 5, 10 and Methods). In contrast, the OCT4 ChIP-seq signal on 50-80 bp subnucleosomes is high at most enhancers (**Fig. 5**, Extended Data Fig. 5 and 10 and Methods). This data shows that the 50-80 bp subnucleosomes form a genomic template that augments OCT4 occupancy of enhancers.

**Figure 5.**
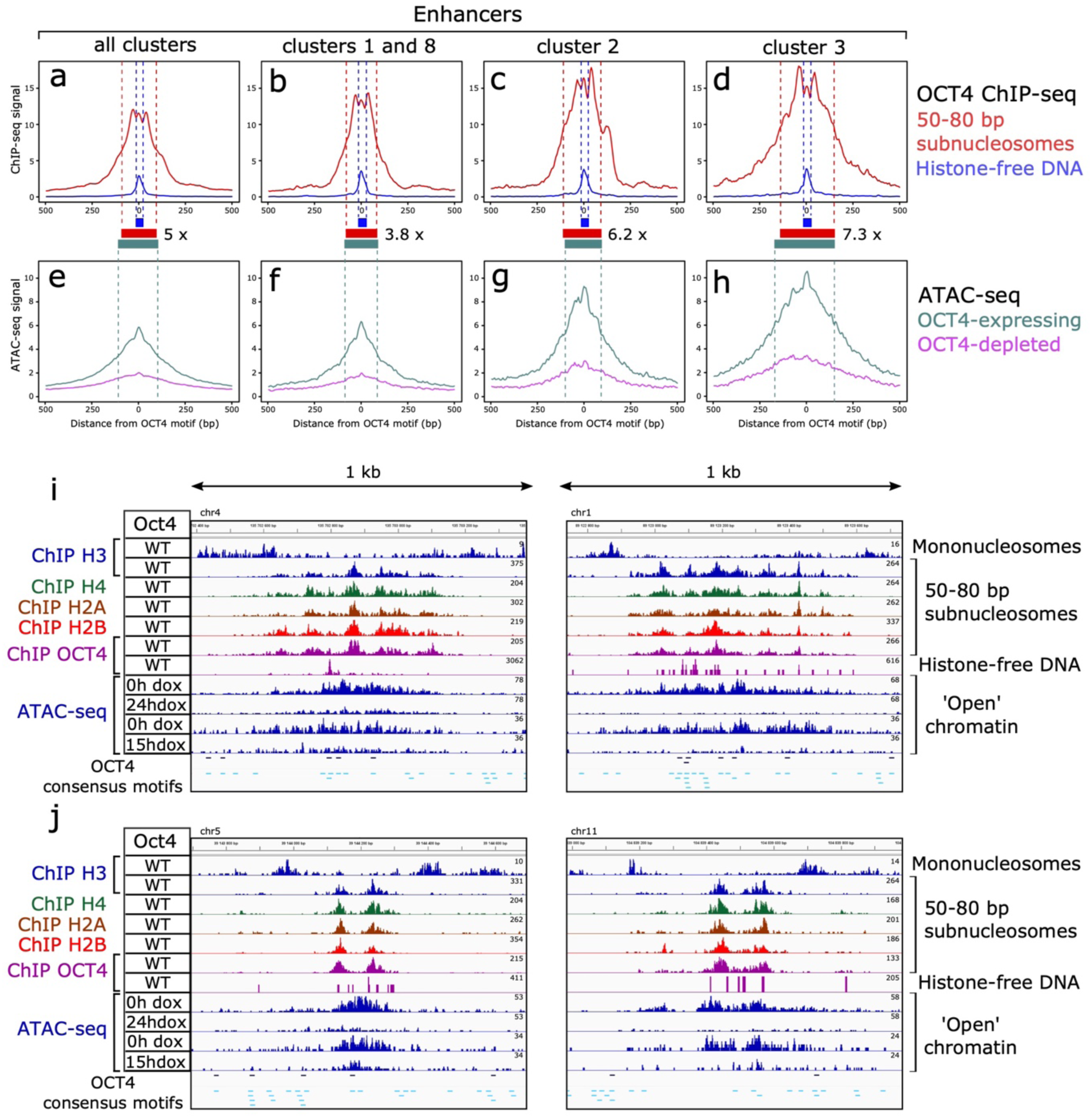
50-80 bp subnucleosomes expand OCT4 binding interval and function in chromatin opening at enhancers. **a**-**d**, Average OCT4 ChIP-seq profiles of 50-80 bp subnucleosomes (red) and histone-free DNA (blue) at enhancers centered on their OCT4 motif. The dashed lines and associated bars delimit the OCT4 enrichment domain by pointing to the positions at which the OCT4 ChIP-seq signal reaches 50% of its maximum. **a**, all enhancers (clusters 1-12 from Extended Data Fig. 1); **b**, single module enhancers (clusters 1 and 8); **c**, **d**, multi-module enhancers of clusters 2 and 3. We selected enhancers containing a single OCT4 high-confidence consensus motif for this analysis (n = 3266, 527, 234 and 230, from **a** to **d,** respectively). Numbers associated with the red bars quantify the expansion of the OCT4 binding interval on 50-80 bp subnucleosomes relative to histone-free DNA. **e**-**h**, Average ATAC-seq profiles of the enhancers selected in the top panels in OCT4-expressing (0h dox) and OCT4-depleted (24h dox) mESCs. The green dashed lines and bar indicate the positions at which the ATAC-seq signal detected in OCT4-expressing cells is at 50% of its maximum. **i**, Density graphs comparing OCT4 ChIP-seq and OCT4-dependent ATAC-seq signals at two representative enhancers of clusters 3. The top lane of each panel shows the positions of canonical nucleosomes detected by histone H3 ChIP-seq of sucrose gradient fractions 11 and 13 in wild-type (WT) mESCs. The next four lanes show the distribution of 50-80 bp subnucleosomes, revealed by histone H3, H4, H2A and H2B ChIP-seq of sucrose gradient fractions 5-6. The following two lanes show OCT4 enrichment detected by ChIP-seq of 50-80 bp subnucleosomes (fractions 5-6) or histone-free DNA from the top gradient fractions (fractions 2-3-4). The next four lanes show the distribution of ATAC-seq signal in OCT4-expressing (0h dox; two independent datasets) or OCT4-depleted (15h or 24h dox) mESCs. High (p = 0.001) and low (p = 0.01) stringency OCT4 consensus binding motifs are indicated in black and blue, respectively. **j**, Two representative enhancers of clusters 1 (left) and 8 (right).

We next hypothesized that the remarkable increase in the size of the OCT4 binding interval on 50-80 bp subnucleosomes might project this TF’s activity beyond the short interval that it binds onto histone-free DNA. Several studies have reported that OCT4 carries a potent function in chromatin opening at enhancers^20,55^. We used the ATAC-seq datasets generated in these studies to test whether OCT4 performs its function in chromatin opening within the genomic interval that it occupies on either histone-free DNA or 50-80 bp subnucleosomes (**Fig. 5e-h**). We observed a striking coincidence of the interval spanned by the OCT4-dependent ATAC-seq signal and the interval occupied by OCT4 on 50-80 bp subnucleosomes (**Fig. 5**). In contrast, the interval occupied by OCT4 on histone-free DNA is much smaller than the extent of the OCT4-dependent ATAC-seq signal (**Fig. 5**). Thus, an essential role of the OCT4-subnucleosome interaction is to expand OCT4 binding interval to project its function in chromatin opening beyond the short region it occupies on histone-free DNA.

## Discussion

The nature of the open chromatin present at eukaryotic cis-regulatory elements has remained elusive due to the difficulty of distinguishing histone-free DNA from fragile nucleosomes and subnucleosomal particles in genome-wide studies. This study used an efficient method to fractionate histone-free DNA fragments, subnucleosomal particles according to their size, and mononucleosomes. We have identified a well-defined organization of open chromatin at enhancers in mouse mESCs, containing five essential components: i) two canonical nucleosomes that flank and delimit the open chromatin region; ii) one or several fragile nucleosome(s); iii) hemisome-like subnucleosomal particles associated to 50-80 bp of DNA, containing the four core histones H2A, H2B, H3 and H4; iv) subnucleosomal particles associated to 81-110 bp of DNA, which could correspond to hexasomes; v) histone-free DNA fragments bound by TFs.

At enhancers, the 50-80 bp subnucleosomes are always present as pairs in which a median distance of 29 bp separates the two particles. The two particles of the pair are symmetrically distributed on each side of the dyad axis of a fragile nucleosome. We have classified enhancers into two categories according to the number of fragile nucleosomes and 50-80 bp subnucleosomes. Single-module enhancers contain one fragile nucleosome associated with two 50-80 bp subnucleosomes, whereas multi-module enhancers contain several fragile nucleosomes and their linked pairs of subnucleosomes. We have shown that this modular organization of enhancers is also observed at super-enhancers and is conserved in human cells.

We have demonstrated that BRG1 is required to generate the 50-80 bp subnucleosomes at enhancers, thus identifying a previously unknown *in vivo* function of this remodeler. By investigating how TFs controlling mESCs pluripotency interact with enhancers, we observed that their highest ChIP-seq enrichment signal coincided with the 50-80 bp subnucleosomes. This binding pattern suggested that the subnucleosomes might form a genuine genomic binding target for TFs at CREs. To test this hypothesis, we compared how the master TF OCT4 interacts with 50-80 bp subnucleosomes and histone-free DNA. As expected, we detected the binding of OCT4 to histone-free DNA at the level of its consensus DNA binding motifs; however, we also found that OCT4 interacts with 50-80 subnucleosomes independently of an OCT4 motif within the particle. We have shown that this absence of OCT4 motifs was not compensated by the presence of degenerated motifs or by half-sites that OCT4 can target onto canonical nucleosomes^11,12^. This result suggests that OCT4 binds a histone domain accessible on the 50-80 subnucleosomes but not on the canonical nucleosomes. Our data supports a model in which BRG1 converts its target fragile nucleosome into two half-nucleosomes (or hemisomes). Each hemisome exposes to the solvent a new histone domain that was hidden before the nucleosome splitting event (**Fig. 4f**). We hypothesize that TFs such as OCT4 recognize and interact with this histone domain to increase their interaction with the open chromatin region of each enhancer.

Our study reveals that promoters and CTCF-BS also exhibit positioned 50-80 bp subnucleosomes within their open chromatin region. Whether these particles might also interact with TFs independently of their DNA binding motif remains to be determined. Such a subnucleosome-associated recruitment pathway could benefit mammalian promoters, which do not contain well-conserved DNA-binding motifs for the general TFs and Pol II^56^. The situation is distinct at CTCF-BS, which include a well-defined consensus motif. Moreover, CTCF occupies most of the DNA molecules containing its binding motif^3^. The subnucleosomes are thus unlikely to be involved in CTCF recruitment. They might instead allow the recruitment of other TFs or proteins that could modulate the function of CTCF-BS.

At the level of enhancers, the interaction of OCT4 with 50-80 subnucleosomes expands the genomic interval occupied by this TF by up to one order of magnitude, compared to the smaller interval it occupies on histone-free DNA (**Fig. 5**). This interaction allows OCT4 to cover the entire length of each enhancer independently of the number and distribution of OCT4 motifs on the DNA. A second function of the OCT4-subnucleosome interaction is to increase OCT4 occupancy. Single-molecule analysis of OCT4 interaction with histone-free DNA has established the low occupancy of its binding motifs in mESCs^3^. In agreement with this previous study, we noticed that only 10% of the enhancers have a robust OCT4 ChIP-seq signal on histone-free DNA. In contrast, the 50-80 subnucleosomes display a strong OCT4 enrichment signal at most enhancers, indicating that these particles increase OCT4 occupancy.

The subnucleosome-mediated expansion of the OCT4-bound interval might benefit enhancer activity by projecting OCT4’s function beyond the boundaries of its occupied DNA motifs. To test this hypothesis, we compared the distribution of OCT4-controlled open chromatin regions with the binding profiles of OCT4 on either histone-free DNA or on the 50-80 bp subnucleosomes. This analysis revealed that OCT4’s potent function in chromatin opening is active precisely within the genomic interval delimited by the interaction between OCT4 and the 50-80 bp subnucleosomes (**Fig. 5**). This domain of OCT4 activity largely exceeds the size of the interval it occupies on histone-free DNA. Our work has thus revealed a novel epigenetic mechanism based on BRG1-generated subnucleosomes that interact with OCT4 to expand its binding interval and project its function beyond the boundaries of its DNA motifs.

## FIGURES

**Extended Data Figure 1.**
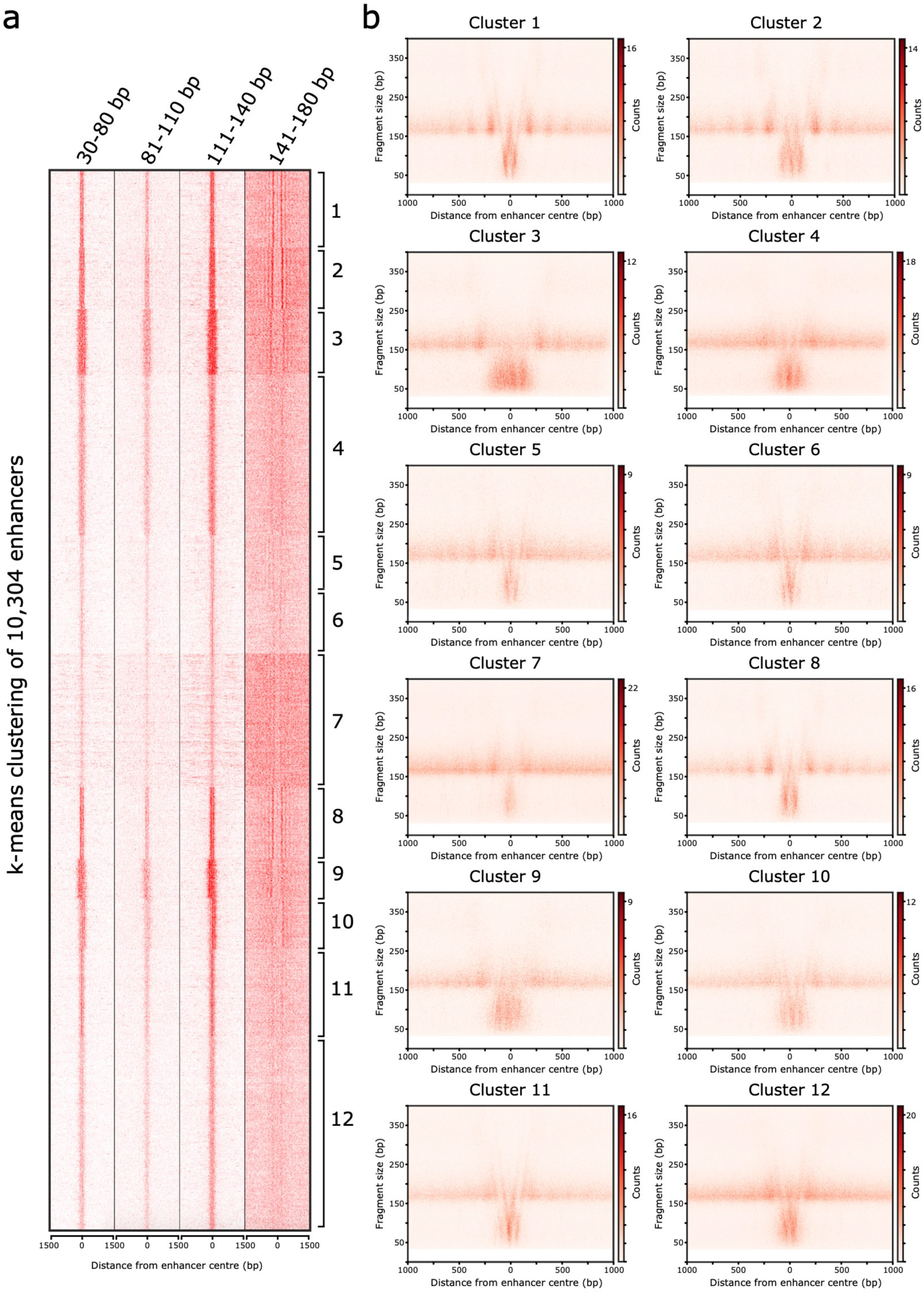
Integrated nucleosomal and subnucleosomal organization at enhancers. **a**, Heat map of histone H3 enrichment at 10,304 enhancers determined by MNase-ChIP-seq (left panel). The four columns display histone H3-immunoprecipitated DNA fragments of different size categories. Enhancers were separated into 12 groups by k-means clustering. The enhancer center corresponds to the middle of its DNase I hypersensitivity peak. **b**, For each group, the distribution of histone H3 ChIP-seq fragments was analyzed by two-dimensional maps (V-plot), showing the distribution and density of DNA fragments immunoprecipitated with antibodies as a function of the position of their mid-point on the genome (x-axis), and their length (y-axis). The color scale corresponds to the number of DNA fragments.

**Extended Data Figure 2.**
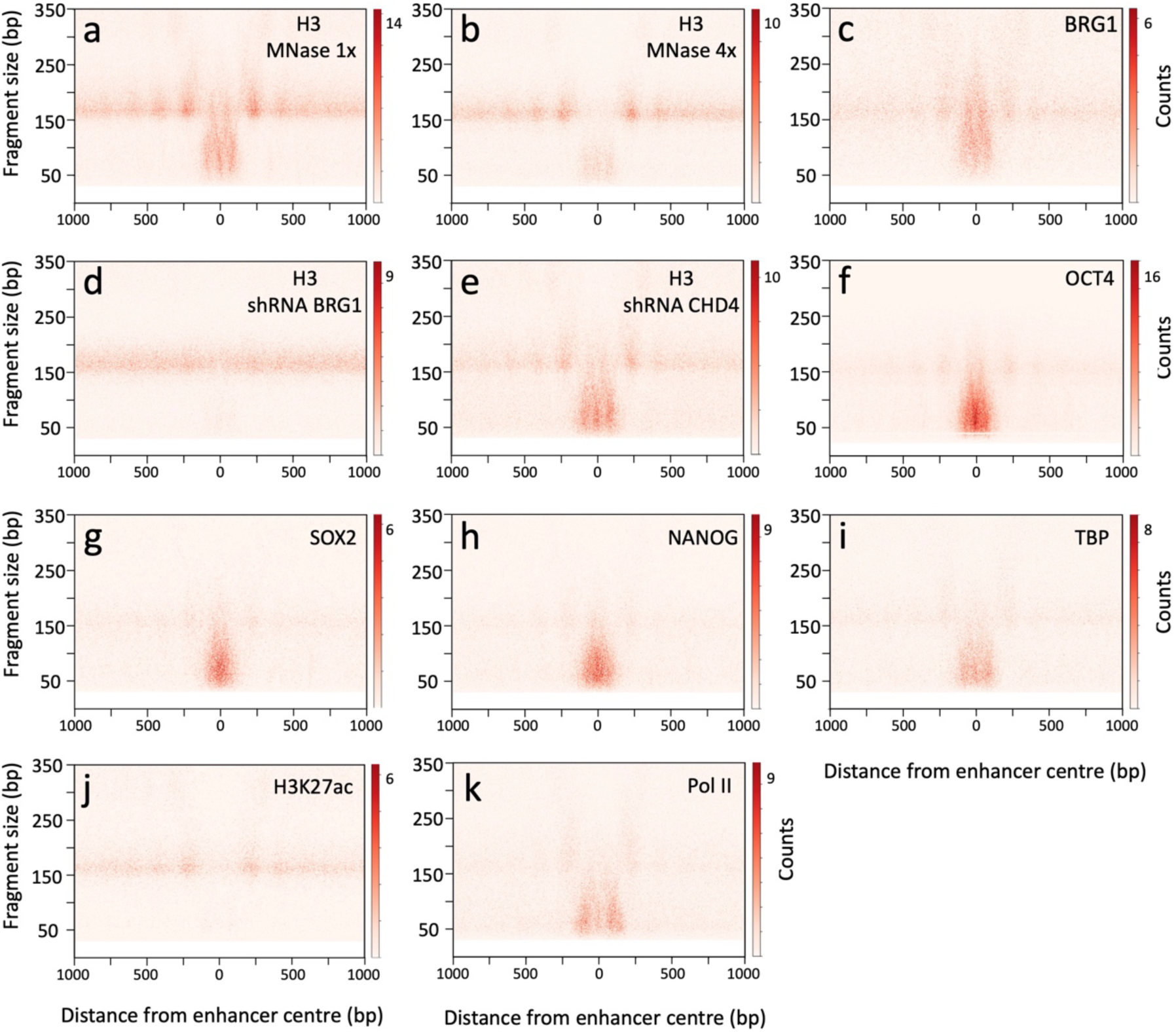
Nucleosomal and subnucleosomal organization of cluster 2 enhancers. **a-b**, V-plots of histone H3 ChIP-seq fragments spanning ± 1000 bp from the enhancer center, using either the standard MNase dose (**a**), or a four-fold excess (**b**). **c**, V-plot of BRG1 ChIP-seq. **d**-**e**, V-plots of histone H3 ChIP-seq in mESCs depleted of either BRG1 (**d**) or CHD4 (**e**), using shRNAs. **f**-**k**, V-plots of OCT4, SOX2, NANOG, TBP, H3K27ac and Pol II ChIP-seq. The standard MNase dose was used in all panels except in (**b**).

**Extended Data Figure 3.**
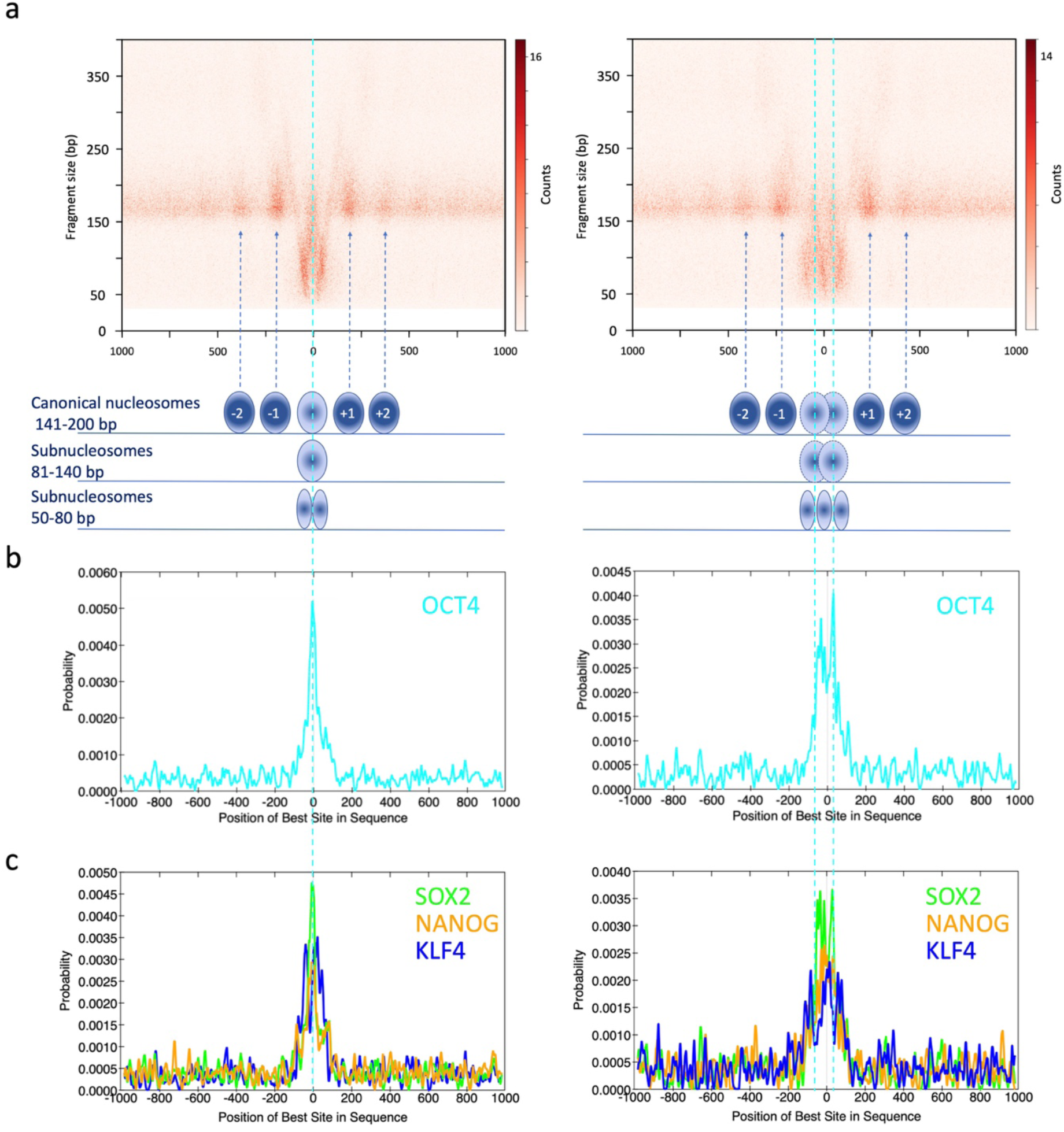
Modular nucleosomal and subnucleosomal organization of enhancers in mESCs. Schematic representations of nucleosomes and subnucleosomal particles are shown below the V-plots of histone H3 ChIP-seq fragments spanning ± 1000 bp from the enhancer center. **a**, Cluster 1 enhancers (left panel) are composed of a single basic module, defined by one fragile nucleosome at the center and two subnucleosomal particles occupying the same genomic interval as the fragile nucleosome. Enhancers from cluster 2 (right panel) are composed of two basic modules, including two fragile nucleosomes in the central region between nucleosomes −1 and +1. These two fragile nucleosomes are separated by about 85 bp and thus cannot coexist on the same molecule. Each of the two fragile nucleosomes is associated with two subnucleosomal particles occupying the same genomic interval. Due to the proximity between the two modules, their subnucleosomal particles partially overlap. (**b, c**) Enrichment for OCT4, SOX2, NANOG and KLF4 consensus BS, defined by centrimo software, within the same genomic intervals as in (**a**).

**Extended Data Figure 4.**
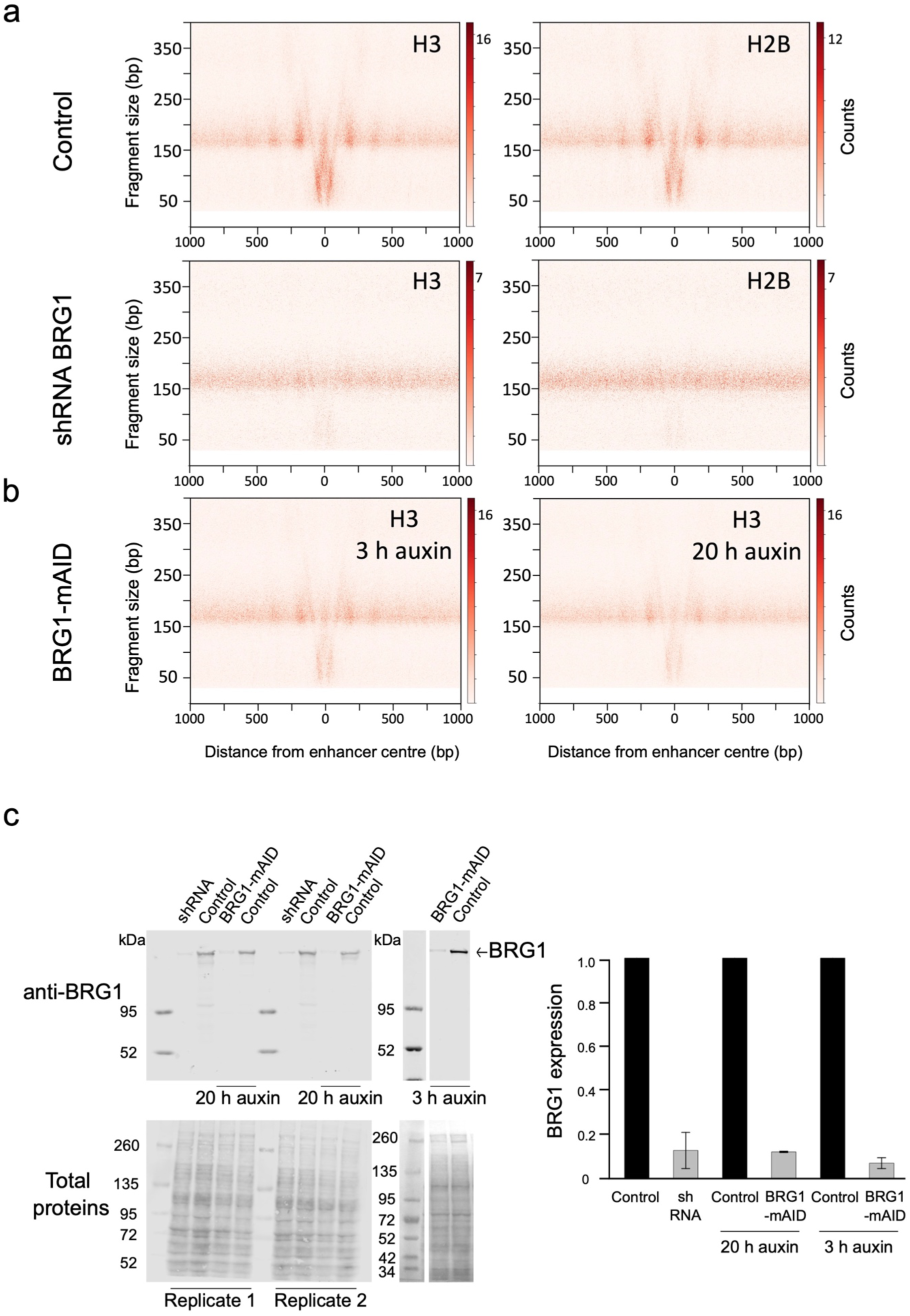
BRG1 depletion induces a reduction in subnucleosomal particles at enhancers and a perturbation of nucleosome positioning. V-plots of ChIP-seq fragments spanning ± 1000 bp from the center of cluster 1 enhancers. **a**, H3 and H2B ChIP-seq were performed on MNase-digested chromatin prepared from mESCs expressing wild-type BRG1 (top panels) or after shRNA-mediated depletion of BRG1 (bottom panels). For the H3 ChIP-seq, depletion of BRG1 was obtained using an shRNA distinct from the shRNA used in Fig. 1. **b**, H3 ChIP-seq was performed using chromatin prepared from mESCs depleted of BRG1 using the AID system by either 3h or 20h of treatment with 1 mM auxin. **c**, Western blot analysis of protein extracts obtained from mESCs transfected with a plasmid expressing an shRNA targeting BRG1, from mESCs depleted of BRG1 using the AID system for either 20 or 3 h or control cells. Ponceau staining of the membrane is shown as a loading control. The right panel shows the means ± sd of BRG1 protein levels in replicate experiments (n shRNA = 8, n 20h auxin = 3 and n 3h auxin = 4).

**Extended Data Figure 5.**
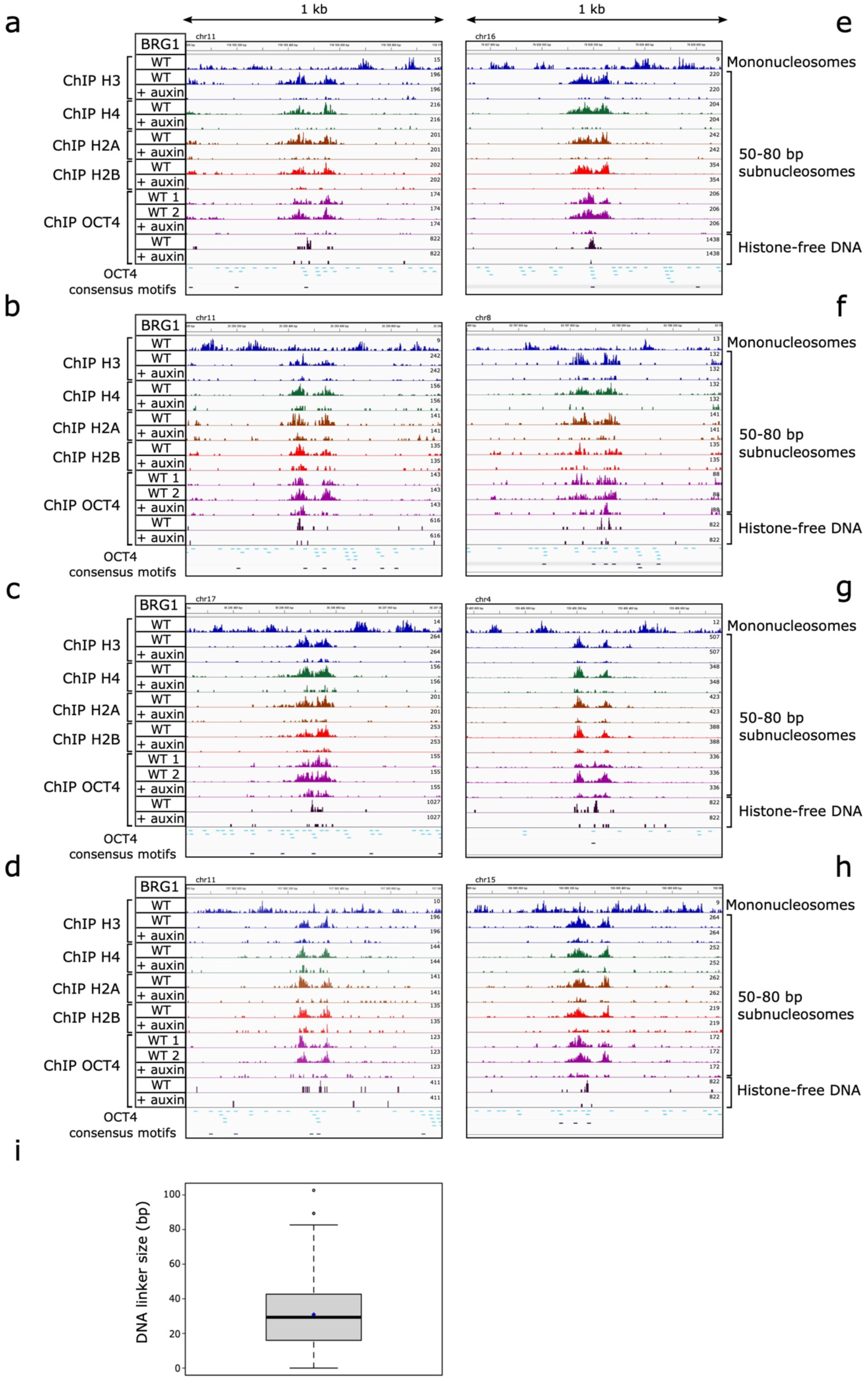
Single-module enhancers accommodate one pair of 50-80 bp subnucleosomes. **a**-**h**, Density graphs showing the distribution of ChIP DNA fragment centers at representative examples of enhancers from clusters 1 and 8. The top lane of each panel shows the positions of canonical nucleosomes detected by histone H3 ChIP-seq of sucrose gradient fractions 11 and 13. The next eight lanes show the distribution of 50-80 bp subnucleosomes, revealed by histone H3, H4, H2A and H2B ChIP-seq of sucrose gradient fractions 5-6. The following five lanes show OCT4 enrichment detected by ChIP-seq of 50-80 bp subnucleosomes (fractions 5-6) or histone-free DNA from the top gradient fractions (fractions 2-3-4). Experiments were performed with chromatin prepared from mESCs expressing wild-type (WT) BRG1 and from cells depleted of BRG1 using the AID system (+ auxin). Two distinct antibodies against OCT4 were used in lanes WT 1 and 2. High (p = 0.001) and low (p = 0.01) stringency OCT4 consensus binding motifs are indicated in black and blue, respectively. **i**, Size distribution of the DNA linker that separates the two 50-80 bp subnucleosomes in single-module enhancers (n = 496, median = 29 bp). The box ranges from the first to the third quartile, the line across the box indicates the median, the blue diamond the mean, the whiskers show the maximum and minimum values of the distribution, and open circles represent outliers.

**Extended Data Fig. 6.**
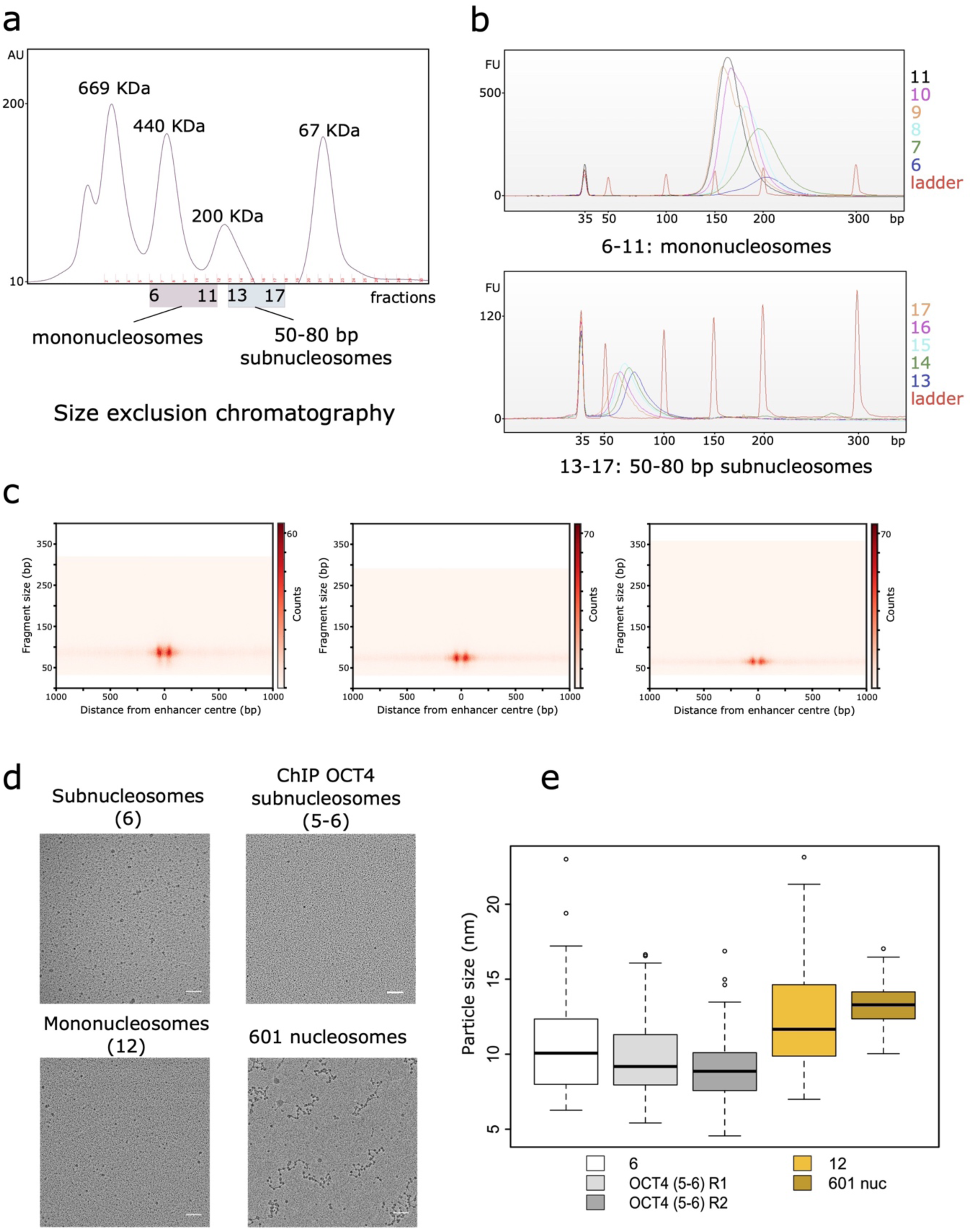
Size analysis of 50-80 bp subnucleosomes. **a**-**c**, Size exclusion chromatography. 50-80 subnucleosomes (sucrose gradient fraction 6) and mononucleosomes (sucrose gradient fraction 11) were independently fractionated onto a Superdex 200 column. **a**, Elution of the mononucleosomes (Superdex 200 fractions 6-11) occurs before that of 50-80 subnucleosomes (Superdex 200 fractions 13-17). The elution of protein size markers is shown for comparison. **b**, The DNA prepared from the mononucleosomes (top) and the 50-80 bp subnucleosomes (bottom) eluted from the Superdex 200 column were analyzed by high-sensitivity DNA electrophoresis. Numbers on the right side of the panels indicate the Superdex 200 fractions. **c**, V-plot spanning ± 1000 bp from the cluster 1 enhancer center of the subnucleosomal particles purified in three consecutive column fractions. **d**, **e**, EM analysis of 50-80 bp subnucleosomes. **d**, The top panels show 50-80 bp subnucleosomes from sucrose gradient fraction 6 and particles from a pool of fractions 5-6 immunopurified with OCT4 antibodies. The bottom panels display mononucleosomes from sucrose fraction 12 and nucleosomes assembled *in vitro* on 601 DNA arrays. **e**, Boxplot showing particle size distribution in the fractions indicated in (**d**). The line across each box indicates the median, boxes indicate the first and third quartile, whiskers the distribution’s maximum and minimum values, and open circles outliers. Two independent OCT4 immunopurification experiments (R1 and R2) were analyzed.

**Extended Data Figure 7.**
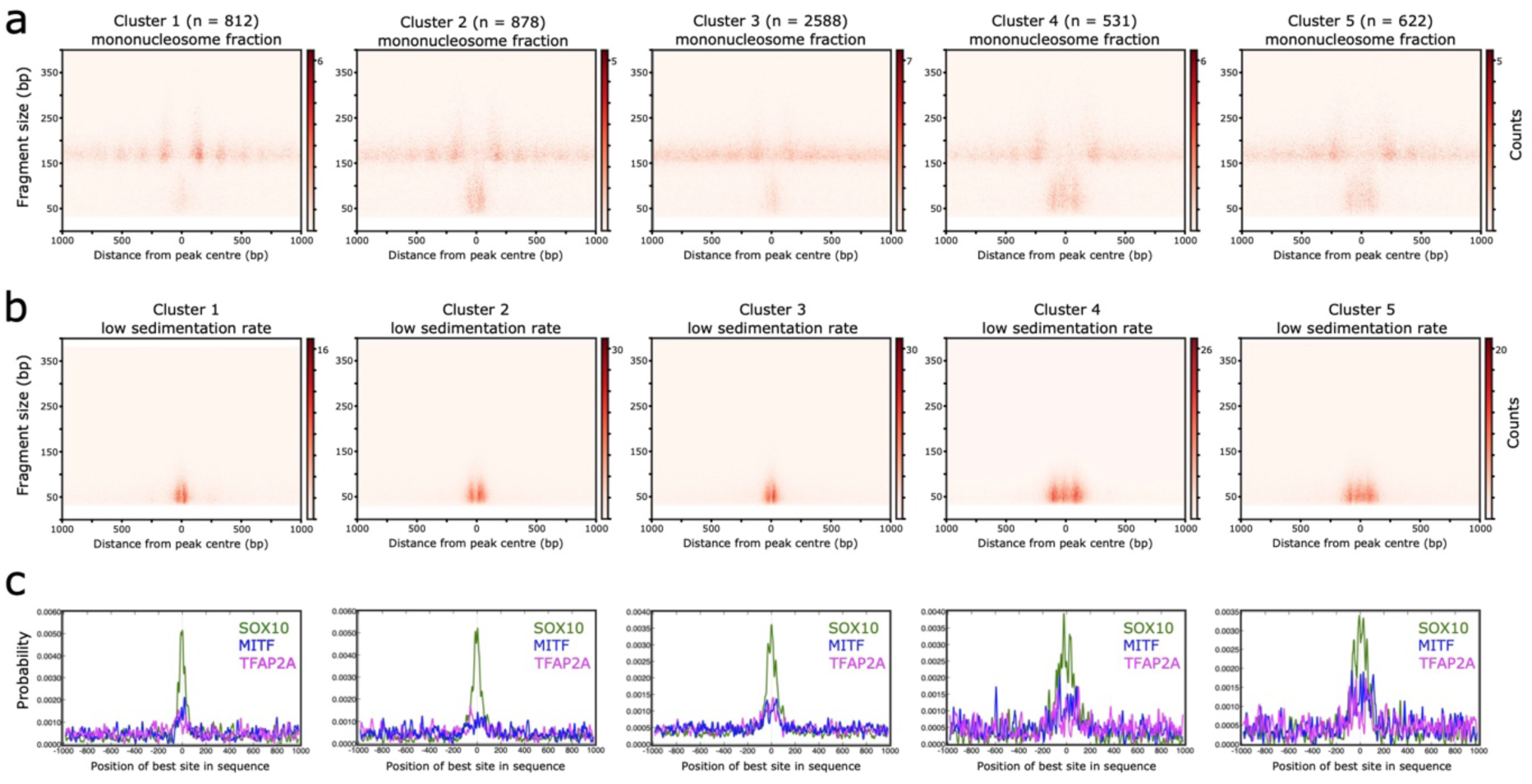
The subnucleosomal organization of enhancers is conserved in human cells. MNase-digested chromatin prepared from human melanoma 501 Mel cells was centrifugated through a 10-30% sucrose gradient, as in Fig. 2. We carried out histone H3 ChIP-seq experiments with gradient fractions containing either mononucleosomes or 50-80 bp subnucleosomes. We used these ChIP-seq datasets for k-means clustering of putative enhancers based on nucleosome and subnucleosome distribution patterns. We isolated five clusters of putative enhancers. **a**, **b**, V-plots of histone H3 ChIP-seq experiments performed with gradient fractions containing mononucleosomes (**a**) or 50-80 bp subnucleosomes (**b**), spanning ± 1000 bp from the center of putative enhancers from each of the five clusters. **c**, Enrichment for SOX10, MITF and TFAP2A consensus BS, defined by Centrimo software, within the same genomic intervals as in (**a**, **b**).

**Extended Data Figure 8.**
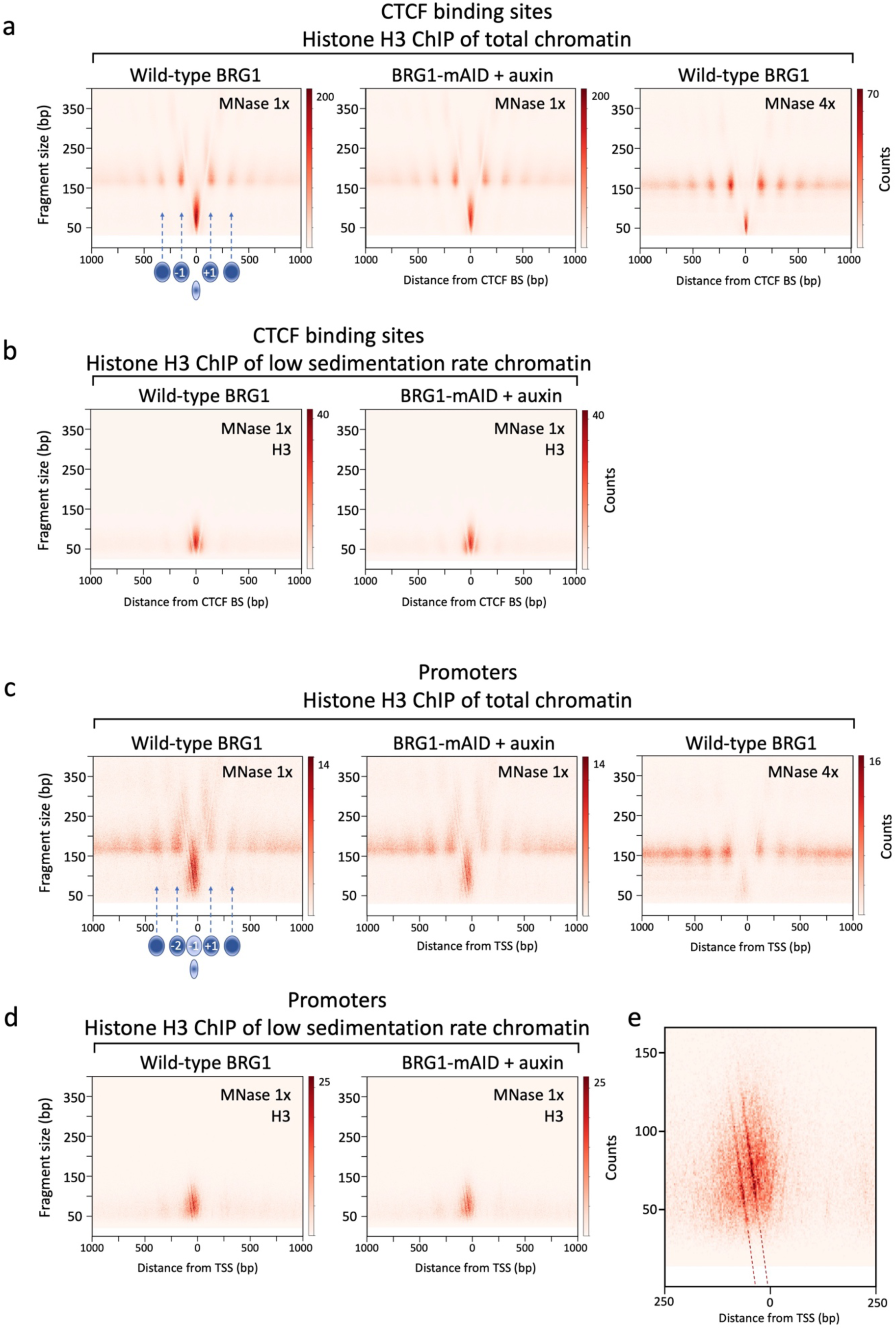
Nucleosomal and subnucleosomal organization at promoters and CTCF-BS. **a**, **c**, V-plots of histone H3 ChIP-seq fragments spanning ± 1000 bp from CTCF-BS (**a**) and TSSs (**c**), using chromatin prepared from mESCs expressing wild-type BRG1 or from cells depleted of BRG1 using the AID system. Chromatin digestion was performed with the standard MNase dose (left and mid panels) or a four-fold excess (right panel). **b**, **d**, MNase-fragmented chromatin, prepared from mESCs expressing wild-type BRG1, or from cells depleted of BRG1 using the AID system, was centrifugated through a sucrose gradient as in Fig. 2. V-plots of histone H3 ChIP-seq experiments performed with sucrose gradient fractions 5-6 reveal 50-80 bp subnucleosomes at CTCF-BS (**b**) and promoters (**d**). The schematic illustrations in (**a**) and (**c**) indicate the positions of nucleosomes and subnucleosomal particles. **e**, Enlargement of **d** (left panel). The two left diagonals result from fragments cleaved precisely on the left and right sides of a region protected from MNase digestion. Extrapolation of the diagonals to y = 0 identifies this region as the −35 to −5 bp interval relative to the TSS. Low molecular weight components of the preinitiation complex might confer this protection.

**Extended Data Figure 9.**
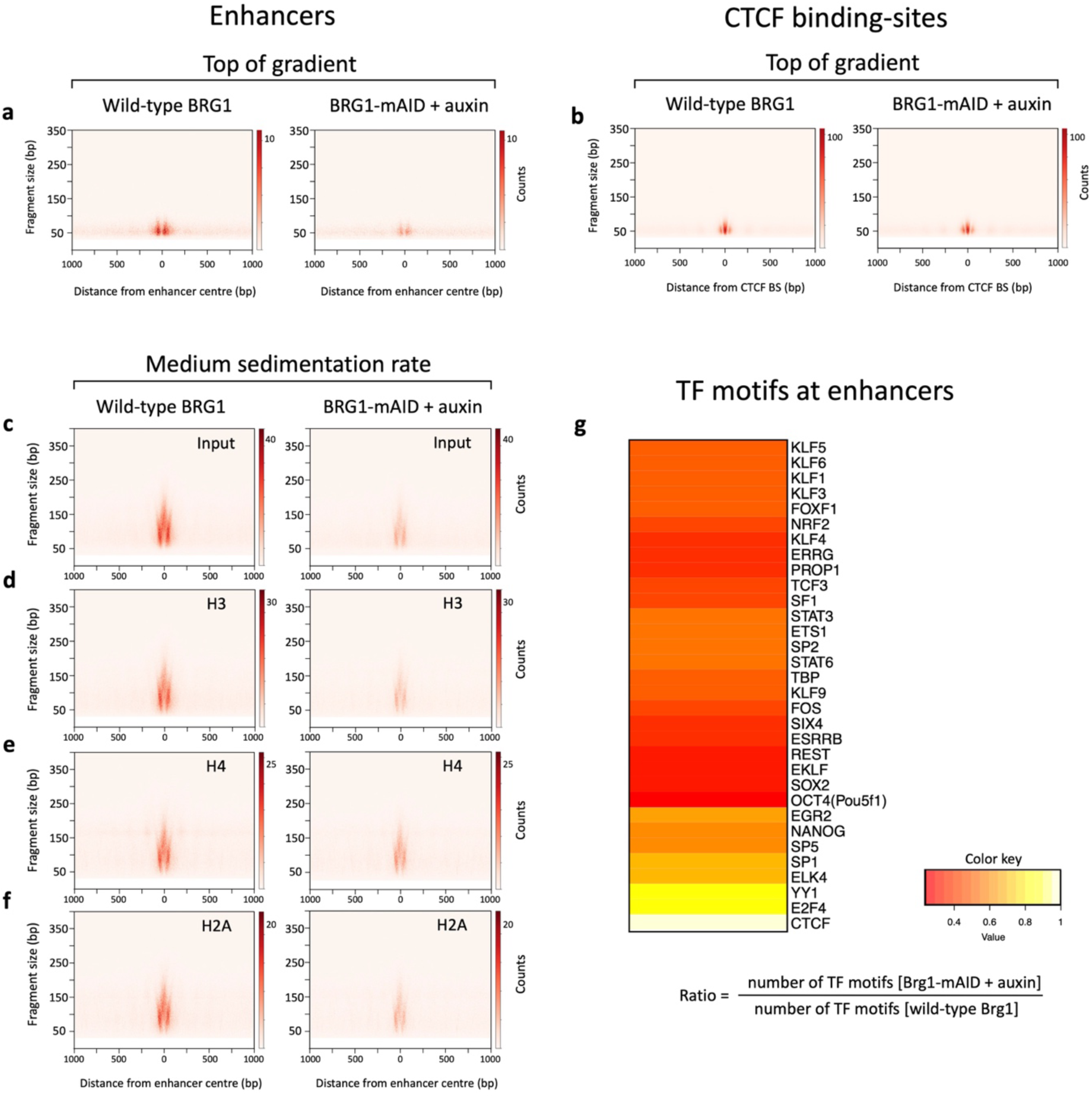
BRG1 depletion alters the production of histone-free DNA and medium sedimentation rate subnucleosomes at enhancers. Chromatin, prepared from mESCs expressing wild-type BRG1, or from cells depleted of BRG1 using the AID system, was centrifugated through a sucrose gradient as in Fig. 2. **a**, **b** V-plots of DNA fragments purified from the top gradient fractions spanning ± 1000 bp from the enhancer cluster 1 center **(a**), or CTCF-BS (**b**). **c**, V-plots of DNA fragments purified from medium sedimentation rate sucrose gradient fractions (pool of fractions 7-8-9) spanning ± 1000 bp from enhancer cluster 1 center. **d**-**f**, V-plots of histone H3, H4 and H2A ChIP-seq experiments performed with the pool of medium sedimentation rate fractions indicated in (**c**) as input. The color scales of each pair of panels were adjusted according to the total number of DNA fragments obtained in each dataset. **g**, TF motif detection with the HOMER tool on the DNA fragments revealed in (**a**). The ratio between the number of TF motifs detected in BRG1-depleted versus BRG1-expressing samples was calculated and visualized as a heatmap. The number of CTCF motifs, which was invariant, was used for normalization.

**Extended Data Figure 10.**
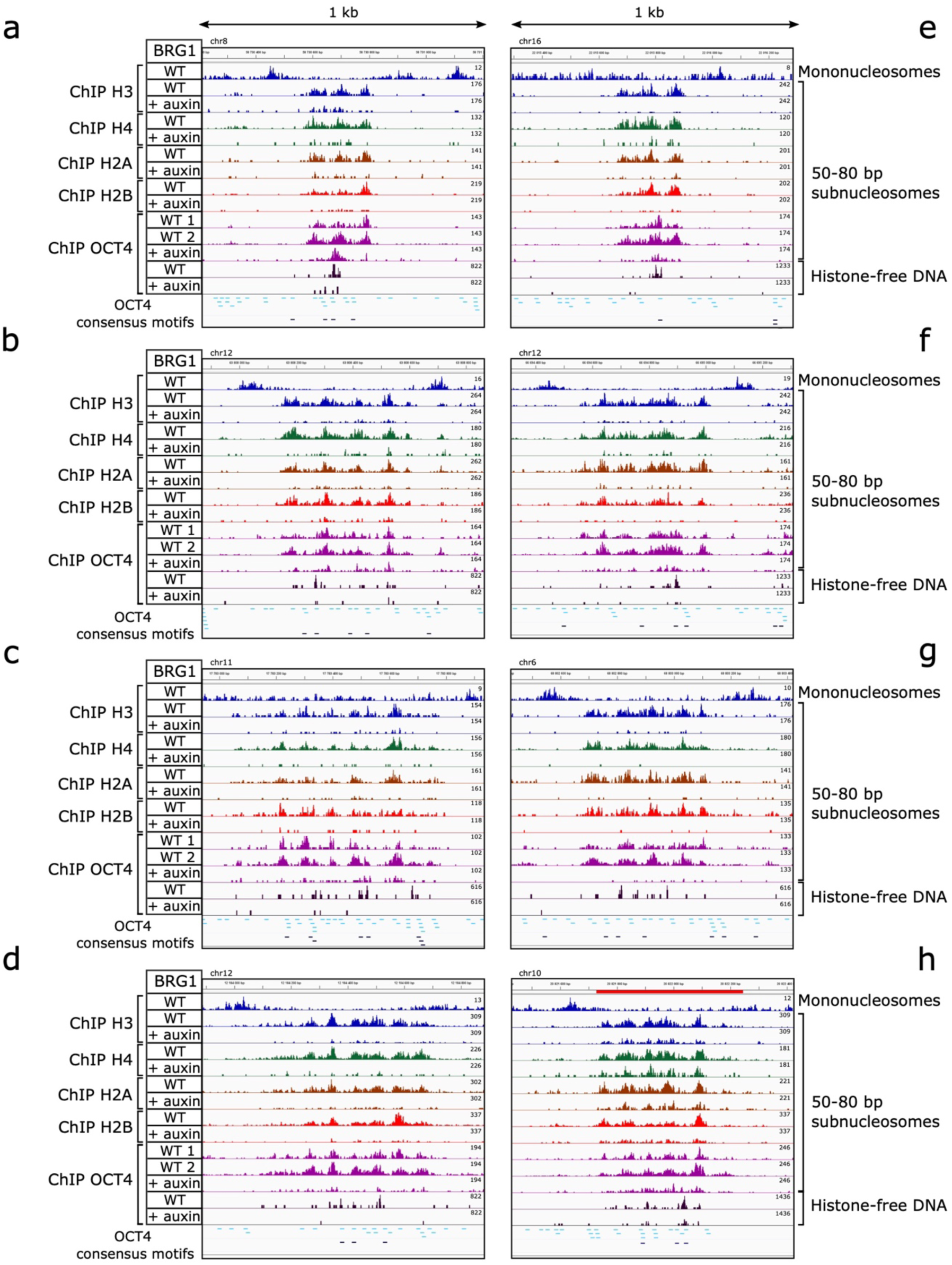
OCT4 enrichment at enhancers coincides with the position of 50-80 bp subnucleosomes. **a**-**g**, Density graphs showing the distribution of ChIP DNA fragment centers at examples of multi-module enhancers from clusters 2 and 3. **h**, Example of an enhancer present within a super-enhancer, showing an organization similar to multi-module enhancers. The top lane of each panel shows the positions of canonical nucleosomes detected by histone H3 ChIP-seq of sucrose gradient fractions 11 and 13. The next eight lanes show the distribution of 50-80 bp subnucleosomes, revealed by histone H3, H4, H2A and H2B ChIP-seq of sucrose gradient fractions 5-6. The following five lanes show OCT4 enrichment detected by ChIP-seq of 50-80 bp subnucleosomes (fractions 5-6) or histone-free DNA from the top gradient fractions (fractions 2-3-4). Experiments were performed with chromatin prepared from mESCs expressing wild-type (WT) BRG1 and from cells depleted of BRG1 using the AID system (+ auxin). Two distinct antibodies against OCT4 were used in lanes WT 1 and 2. High (p = 0.001) and low (p = 0.01) stringency OCT4 consensus binding motifs are indicated in black and blue, respectively.

**Extended Data Figure 11.**
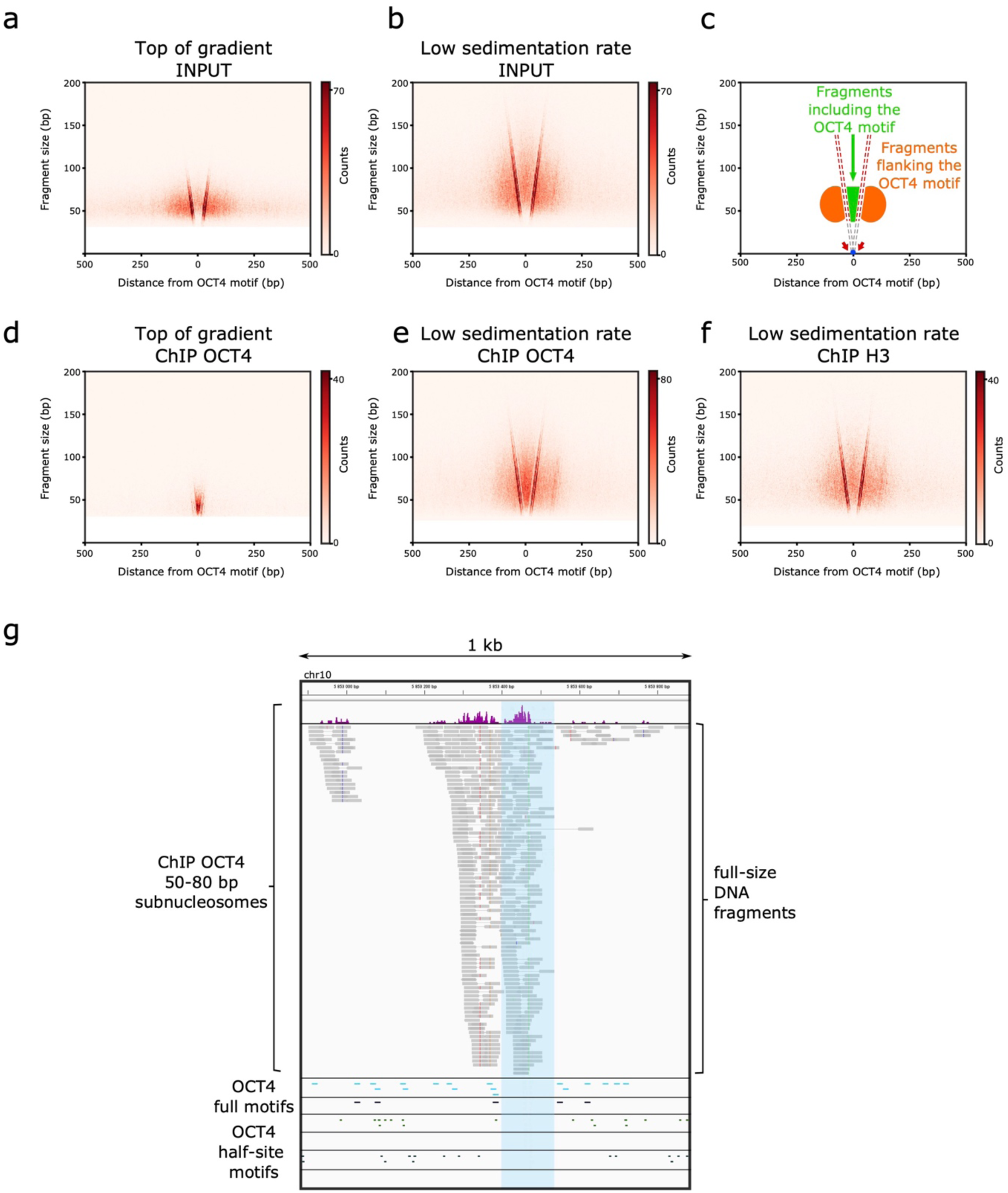
OCT4 interacts with 50-80 bp subnucleosomes lacking its consensus binding motif. **a**-**b** and **d**-**f**, V-plots of enhancers centered on their OCT4 motif. Enhancers containing a single OCT4 consensus binding motif were selected for this analysis (n = 1,302). MNase-fragmented chromatin, prepared from mESCs expressing wild-type BRG1, was centrifugated through a sucrose gradient as in Fig. 2. **a**-**b**, V-plots of DNA fragments purified from the top gradient (**a**) and low sedimentation rate (**b**) fractions. **c**, Schematic diagram illustrating key features of V-plots centered on OCT4 motifs. The green area corresponds to MNase-released DNA fragments containing the central OCT4 consensus binding motif. The two orange regions are defined by adjacent or proximal DNA fragments that do not include the OCT4 motif. The red arrows indicate MNase hypersensitive sites on each side of the OCT4 motif, which are at the origin of the diagonals forming the ‘V’ on each map. **d**-**e**, Maps of OCT4 ChIP-seq experiments of the top gradient (**d**) and low sedimentation rate gradient (**e**) fractions, using as input the fractions shown in (**a**, **b**). **f**, Map of histone H3 ChIP-seq experiment of low sedimentation rate gradient fractions. **g**, Example of a single-module enhancer. The top lane shows in purple the density graph of the DNA fragment centers from OCT4 ChIP-seq 50-80 bp subnucleosomes. The genomic position of each individually sequenced fragment is indicated below in grey. Blue shading highlights the DNA fragments that do not contain OCT4 motifs. High (p = 0.001) and low (p = 0.01) stringency OCT4 consensus binding motifs are indicated in black and blue, respectively. The next four lanes show in green the positions of low (p = 0.01) and high (p = 0.001) stringency OCT4 POU_S_ (top two lanes) and POU_HD_ (bottom) half-sites; no half-site motif was detected at high stringency in the 1 kb interval.

## Methods

### Cell culture

mESCs (46C, E14Tg2a and their derivatives) were grown on mitomycin C-inactivated mouse embryonic fibroblasts at 37 °C, 5 % CO_2_, in DMEM (Sigma) supplemented by leukaemia inhibitory factor (LIF), 1X non-essential amino acids (Invitrogen), 15 % fetal calf serum (Invitrogen), 0.2 % β-mercaptoethanol (Sigma) and 1X penicillin/streptomycin (Invitrogen)^57^. The human melanoma 501Mel cell line^58^ was grown in RPMI 1640 medium (Sigma) supplemented with 10% fetal calf serum and 1X penicillin/streptomycin.

### Plasmid construction

We designed the shRNA-expressing plasmids as previously described^59^. Sequences of the sense strand of shRNAs targeting BRG1 are as follows: shRNA O5: 5’-GCTCCAGTAAAGATGTCTACT-3’; shRNA O7: 5’-GAGCGAATGCGGAGGCTTATG-3’. Sequences targeting *Chd4* were already described^18^.

The AID system is based on the fusion, at the C-terminus of the target protein, of a mini AID (mAID) tag (68 amino-acids, 7.4 kDa)^41^. We adopted the two-selection marker strategy to target the *Smarca4* alleles with the sequence encoding mAID^41^. We assembled the *Smarca4*-mAID targeting vector by a serial modification of the base vectors pMK287 (mAID-Hygro, Addgene #72825) and pL452^18^. First, we introduced two adapters, upstream and downstream of the region coding for mAID and Hygromycin resistance into the pMK287 vector. The upstream adapter contained a sequence encoding a (GGGGS)_3_ linker in front of the mAID coding sequence. The region containing the linker-mAID sequence was then amplified by PCR using Phusion polymerase (NEB) from the modified pMK287-mAID-Hygro plasmid and subcloned into the pL452 vector which harbors the Neomycin resistance cassette under the control of a PGK promoter. We amplified the left and right homology arms (400-500 bp each) corresponding to the last exon and 3’UTR of *Smarca4* by PCR from E14Tg2A genomic DNA. We designed the oligonucleotides that amplify homology arms to introduce silent mutations after integrating the mAID cassette downstream of the *Smarca4* gene, to prevent re-cutting by the Cas9 enzyme. In the next cloning steps, we introduced the PCR products of left and right homology arms using restriction enzyme digestions into both vectors (pMK287-mAID-Hygro and pL452-mAID-Neo).

We assembled the CRISPR/Cas9 vectors by annealing pairs of oligos carrying the sgRNA sequences and cloning them into pSpCas9(BB)-2A-Puro (PX459) (Addgene #62988) as described^60^. Two distinct *Smarca4*-targeting sgRNAs were cloned separately by annealing oligos caccgTTGGCTGGGACGAGCGCCTC and aaacGAGGCGCTCGTCCCAGCCAAc for the first sgRNA and caccgGCGCCTCGGGGTCAGGACTC and aaacGAGTCCTGACCCCGAGGCGCc for the second sgRNA. We designed the sgRNAs to target the area close to the last exon of *Smarca4* using the Zhang lab guide design tool (https://zlab.bio/guide-design-resources).

### Generation of the Smarca4-mAID cell line

We used the E14Tg2a-Tir1 cell line, which was obtained by targeting the *Tigre* locus of E14Tg2a cells with a transgene expressing *Os* Tir1^61^. Using the Neon electroporation kit (ThermoFisher), we transfected this cell line with the two vectors encoding the Cas9 enzyme and sgRNAs (2.5 μg each) and with the two linearized plasmids carrying the *Smarca4* homology arms and the mAID tag (pMK287, pL452, 10 μg each). Neomycin (250 μg/ml, G-418, Sigma) and Hygromycin B (300 μg/ml, Thermo Fisher Scientific) selection was started after 24 h. After six days of selection, single clones were manually picked, expanded in 24-well plates and subjected to genotyping by PCR and sequencing of the knock-in region. Western blotting of protein extracts from auxin-treated cells confirmed selected clones with a homozygous insertion of the mAID cassette at the *Smarca4* locus.

### Chromatin remodeler depletion

For shRNA-mediated depletion, we transfected two million mESCs by electroporation using Amaxa nucleofector (Lonza) or Neon kits, with 15 µg of pHYPER-linker control plasmid or a pHYPER plasmid containing remodeler-specific shRNA. Puromycin (2 µg/ml) was added to the culture medium 24h after electroporation, and the cells were grown under selection for an additional 30h (Chd4 shRNA) or 48h (BRG1 shRNA) period, as previously described^18^. Western blot experiments validated the efficiency of remodeler depletion.

For auxin-mediated protein depletion, we supplemented the cell culture medium of cells expressing BRG1-mAID with 1 mM auxin (IAA, indole-3-acetic acid, Cayman Chemical) for 3 h or 20 h. Unless specified otherwise, we realized all auxin-mediated depletion of BRG1 experiments described in the manuscript at the 20h auxin timepoint.

### Chromatin preparation

mESCs were treated with trypsin to obtain a single-cell suspension and separated from mouse embryonic fibroblast feeders by a 30 min incubation at 37°C in D15 medium on a gelatin-coated culture plate. mESCs were collected and fixed in D15 culture medium containing 1 % formaldehyde (Sigma) for 10 min at 20°C. 501Mel cells were treated with trypsin and fixed in culture medium as described for mESCs. We stopped fixation by adding glycine to 125 mM, and the cells were centrifugated and washed three times with PBS. Cells were then permeabilized in Tris-HCl pH 7.5 15 mM, NaCl 15 mM, MgCl_2_ 5 mM, EGTA 0.1 mM, KCl 60 mM, Sucrose 0.3 M, 0.4% IGEPAL CA-630 (I3021, Sigma) during 15 min on ice. We next diluted the cells with two volumes of MNase buffer (Tris-HCl pH 7.5 20 mM, NaCl 20 mM, CaCl_2_ 2 mM, MgCl_2_ 4 mM, KCl 15 mM, 0.1 M sucrose), and chromatin was digested during 10 min at 37 °C with 6 Kunitz units of MNase (New England Biolabs, 200 Kunitz units/µl) per 1 million cells. We stopped MNase digestion by adding a final concentration of 10 mM EDTA. We released chromatin from MNase-treated mESCs by passing the cell suspension 13 times through a 26G syringe. Next, we centrifuged the samples, and the supernatants containing the solubilized chromatin were used directly for ChIP-seq experiments or sucrose gradient centrifugation. The quality and reproducibility of the MNase digestion pattern of all samples were validated after crosslink reversal by electrophoresis on a 1.7 % agarose gel.

### Fractionation of MNase-digested chromatin by sucrose gradient centrifugation

Chromatin samples prepared from 40 million ES or 501 Mel cells were supplemented with protease inhibitors (Complete Protease Inhibitor Cocktail, Roche) and loaded on a 9.9 ml manually poured 10-30 % sucrose gradient containing 10 mM Tris-HCl pH 7.5, 1 mM EDTA, 1 mM EGTA, 80 mM NaCl and protease inhibitors. We centrifuged the chromatin samples for 16 h at 4 °C, 40,000 rpm (197,000 g) using a Beckman Coulter SW 41 swinging-bucket rotor. After centrifugation, 20 chromatin fractions of 500 µl were collected from top to bottom of the tube and stored at −80°C. Aliquots of each chromatin fraction were treated with RNAse A and proteinase K, and subjected to reverse crosslinking for 16 h at 65 °C, followed by phenol-chloroform DNA extraction and ethanol precipitation. DNA was quantified using the Qubit system (Thermo) and analyzed on a 4% agarose gel or used for library preparation and deep sequencing.

### MNase ChIP-seq

We realized all ChIP-seq experiments with formaldehyde-fixed, MNase-fragmented chromatin, either with individual sucrose gradient fractions, pools of gradient fractions, or whole-cell chromatin extracts. For the latter, we used chromatin solubilized from 1.2 million mESCs for ChIP-seq with antibodies against histones or from 10 million cells for ChIP-seq with antibodies against TFs and Pol II. Chromatin samples were incubated with antibodies (20 µg of antibodies against TFs or Pol II and 5 µg of antibodies against histones) overnight at 4 °C. Chromatin-antibody complexes were next incubated with 50 µl of protein G-agarose beads for 4 h at 4 °C. Then, chromatin-antibody-beads complexes were washed three times with TEN buffer (20 mM Tris-HCl pH 7.5, 150 mM NaCl, 3 mM MgCl_2_, 0.1 mM EDTA, 0.01% Igepal), four times with WBLiCl buffer (50 mM Hepes pH 7.5, 500 mM LiCl, 1 mM EDTA, 1 % IGEPAL, 0.7 % Na-deoxycholate) and once with TEN buffer. Between each wash, we centrifuged the beads for 3 min at 800 g, 4 °C. Elution was realized by resuspending the beads in elution buffer (50 mM Tris-HCl pH 8, 10 mM EDTA, 1% SDS). For ChIP-seq experiments realized with the antibody targeting the 20 C-terminal amino acids of OCT4 (Fig. 4c and WT 1 lane of Fig. 4d, Extended Data Fig. 5 and 10), we performed the washes and elution by peptide competition as described below for sequential ChIP-seq of OCT4- and H2B-bound chromatin. Eluted chromatin was treated with proteinase K and subjected to reverse crosslinking and phenol/chloroform extraction, followed by ethanol precipitation. We prepared DNA libraries using MicroPlex Library Preparation Kit v2 or v3 (Diagenode). Sequencing was performed on a NextSeq 500 Illumina platform using NextSeq 500 High Output v2 (75 cycles) kit. At least two biological replicates were realized and sequenced for each histone or TF ChIP-seq experiment.

### Sequential ChIP-seq of OCT4- and H2B-bound genomic DNA

We used chromatin prepared from 240 million E14Tg2a mESCs for each experiment. We incubated the chromatin with either 45 µg antibodies against OCT4 (MBS420786, MyBioSource) or 45 µg of non-specific IgG as a control. After 16 h at 4 °C, we added Protein G-agarose beads to chromatin-antibody complexes and further incubated for 4 h at 4 °C. After seven washes with TEN buffer, we incubated the beads with TEN buffer containing OCT4 peptide (C-terminal part of OCT4 sequence: H-Cys-Lys-Lys-Lys-Lys-Pro-Ser-Val-Pro-Val-Thr-Ala-Leu-Gly-Ser-Pro-Met-His-Ser-Asn-OH) for elution by competition. We performed three consecutive elution steps: 2 h at room temperature with TEN buffer containing 2 mg/ml of OCT4 peptide, 2 h at room temperature with 1 mg/ml of peptide, and 16 h at 4 °C with 1 mg/ml peptide. The three elution fractions were pooled and incubated with 5 µg of antibodies against histone H2B for 16 h at 4 °C. Protein G-agarose beads were next added and incubated for 4 h at 4 °C. We performed the washes and elution steps of this second ChIP as described above for regular ChIP-seq experiments. Two biological replicates were obtained for each OCT4 and non-specific IgG antibody.

### Transmission Electronic Microscopy

Chromatin samples were prepared from 80 million E14Tg2a mESCs and centrifugated through a sucrose gradient, as described above for MNase ChIP-seq. We examined subnucleosomal particles collected from fractions 5, 6, 11 and 12 of the sucrose gradient and particles further purified from a pool of fractions 5-6 by immunoprecipitation with the antibody targeting the 20 C-terminal amino-acids of OCT4, as described above. We analyzed half of each replicate sample of OCT4-immunopurified particles by EM, as shown in Extended Data Fig. 6d,e. We processed the other half as a ChIP-seq sample, which was used for the V-plot shown in Fig. 4c and for the WT1 lane of the density graphs shown in Fig. 4d and Extended Data Fig. 5 and 10.

Aliquots of sucrose gradient fractions were dialyzed overnight into TEAP10 buffer (10 mM Triethanolamine-HCl (pH 7.5), 0.2 mM EDTA, 10 mM NaCl, 0.2 mM PMSF), then crosslinked with 0.1% glutaraldehyde at 4°C for 2 h and quenched by the addition of Tris (pH 8) to a final concentration of 50 mM. Samples were coated with 2 × 10^−4^ % benzalkonium chloride (BAC) for 1 h at room temperature, and 5 µl of the sample (approximately 0.02 ng/ µl) was adsorbed onto carbon/formvar supported 200 mesh copper grids (TAAB technologies). Grids were washed twice with deionized water, dehydrated with 90% ethanol, and blotted dry. Grids were rotary shadowed with 3 nm platinum at a 7° angle and examined on the JEOL JEM-1400Plus Transmission Electron Microscope.

We randomly selected the particles and measured them with Image J for statistical analysis of particle size. Most particles had a disk-like structure, and we measured them along the diameter. We measured particles having an elliptical shape along the major axis, as this likely represents the front view of the nucleosomal particles. We built the boxplot shown in Extended Data Fig. 6e with the sizes of 92, 102, 97, 71 and 100 particles from one sample of sucrose gradient fraction 6, two independent samples of OCT4-immunopurified particles (OCT4 (5-6) R1 and R2), one sample of fraction 12 (mononucleosomes) and one preparation of *in vitro*-assembled 601 nucleosomes, respectively. To choose the appropriate statistical test to compare the different groups of particles, we first verified the size distribution in each group and performed Shapiro-Wilk normality tests. This analysis revealed that the data collected for all groups of subnucleosomal particles did not display a normal distribution. We thus selected a Mann-Whitney test (R Wilcox.test function; two.sided; paired = FALSE; conf.level = 0.95 parameters) to determine if the sizes of subnucleosomal particles and canonical nucleosomes were significantly different. This statistical analysis revealed that subnucleosomal particles of each group are significantly different in size from canonical nucleosomes; particles from fraction 6 versus those from fraction 12: p-value = 6.6 × 10^−05^; particles from OCT4 (5-6) R1 versus those from fraction 12: p-value = 1.5 × 10^−07^; particles from OCT4 (5-6) R2 versus those from fraction 12: p-value = 2.9 × 10^−11^, particles from fraction 6 versus 601 nucleosomes: p-value = 2.4 × 10^−14^; particles from OCT4 (5-6) R1 versus 601 nucleosomes: p-value < 2.2 × 10^−16^; particles from OCT4 (5-6) R2 versus 601 nucleosomes: p-value < 2.2 × 10^−16^.

### Size exclusion chromatography

Chromatin samples were sucrose gradient fraction 6 for 50-80 bp subnucleosomes and fraction 11 for mononucleosomes. 400 µl of chromatin sample were loaded at 0.25 ml/min onto a Superdex 200 increase 10/300 GL column (Cytiva) equilibrated in 10 mM Tris-HCl pH7.5, 80 mM NaCl, and 1 mM EDTA. 250µl fractions were collected. The column was calibrated with gel filtration protein markers: thyroglobulin (669 kDa), apoferritin (440 kDa), b-amylase (200kDa) and bovine serum albumin (67 kDa) from MWGF1000, Sigma).

### Construction of the hemisome-link-hemisome model

To generate the hemisome-linker-hemisome molecular model, we separated the best resolution nucleosome structure (1kx5 bp) into two hemisomes by cutting the DNA at the dyad position. We positioned a B-DNA linker at the generated DNA extremity by overlapping four base pairs. We choose a representative length of 29 bp corresponding to the median distance between two hemisome-like particles at enhancers. The second hemisome was positioned similarly at the second extremity of the DNA linker. The exposed DNA is highly sensitive to MNase and thus digested prior to the ChIP experiment. The resulting isolated particles, corresponding to hemisomes, would contain 50-70 bp DNA fragments linked to histones H3-H4-H2A and H2B.

### Computational analyses

#### Lists of mESCs enhancers

We first mapped the distribution of cis-regulatory elements onto the mESC genome using DNaseI-seq data (GSM1014187), identifying 139,454 DNase-hypersensitive (DHS) regions, as previously described^18^. Gene promoters were filtered out using RefSeq coordinates of TSSs, and putative enhancers were identified based on the co-occurring binding of pluripotency-associated TFs OCT4 (GSM1082340), SOX2 (GSM1082341) and NANOG (GSM1082342), as well as of the mediator (GSM560347)^25,53^. To avoid potential interferences with CTCF-bound regions in our nucleosomal organization analysis, we removed from these putative enhancers those bound by CTCF, using mESCs datasets GSM723015 and GSM1828650. 19,365 loci displaying high ChIP-seq signal for OCT4, SOX2, NANOG, and the Mediator, but negative for CTCF, were identified as putative enhancer elements. We then used the histone H3 MNase ChIP-seq datasets generated in this study to analyze nucleosomal and subnucleosomal organization at these putative enhancers. One of these H3 MNase ChIP-seq datasets was converted into a BED file and subdivided into five files containing DNA fragments of 30-80, 81-110, 111-140, 141-180, and >181 bp. The number of DNA fragments in each subgroup was as follows. 30-80 bp: 13,806,227 DNA fragments; 81-110 bp: 9,480,961 fragments; 111-140 bp: 13,589,276 fragments; 141-180 bp: 64,258,774 fragments; >180 bp: 51,518,836 fragments. Next, in each BED file, DNA fragments were reduced to 10 bp and centered at the level of the fragment midpoint. Using seqMINER^62^, fragment densities from these modified BED files were collected in a −1500/+1500 bp window around the center of each enhancer (defined in this study as the center of the DHS peak) and subjected to several rounds of k-means clustering. This analysis allowed us to classify 10,304 (out of the initial 19,365) putative enhancers into the 12 clusters shown in Extended Data Fig. 1.

We developed the exPATT.R tool to detect OCT4 DNA binding motifs and change the genomic coordinates of each enhancer to have the OCT4 motif at its center. The exPATT.R R script filters genomic regions, recorded in an input BED file, that match a given DNA pattern (OCT4 consensus motif in this study) depicted as a position weight matrix (PWM). These pattern matrices were matched in genomic region using the PWM-related functions of the Bioconductor package Biostrings^63^ with a minimum score of 80% of the highest possible score. Then, genomic regions with at least one pattern hit were filtered. The output of this script was a new BEDGRAPH file with filtered DNA regions of 2 kb centered around the pattern. All BED file input and output operations were performed using the rtracklayer package. Then, we used bedtools to select enhancers bearing a single central OCT4 motif per enhancer (n = 1,302, Extended Data Fig. 11).

#### Lists of 501 Mel cell enhancers

Our extensive analysis of the distribution of 50-80 bp subnucleosomes in mESCs revealed their accumulation at enhancers, promoters and CTCF-BS (this study). Based on the hypothesis that subnucleosomal particles are conserved in human cells, we mapped cis-regulatory elements in human 501 Mel cells using the histone H3 ChIP-seq dataset from 50-80 bp subnucleosomes purified from these cells. We used MACS2 to call 91,205 peaks from this dataset. We next focused our analysis on putative enhancer elements located in intergenic regions. We filtered out all regions of the human genome located within 3 kb of gene TSSs or 10 kb of gene end coordinates. We also set up a strategy to remove CTCF-bound loci from this list of peaks. Since the CTCF ChIP-seq distribution has not been defined in 501 Mel cells, we took advantage of the observation that CTCF binding to mammalian genomes is relatively independent of cell type^64,65^. We removed all loci bound by CTCF in two cancer cell lines from the putative enhancer list, using ENCODE CTCF ChIP-seq datasets GSM749683, GSM749715, GSM749679 and GSM749768. This combination of filtering steps led to a list of 14,502 putative 501 Mel cell enhancers. K-means clustering of these 14,502 elements using the H3 ChIP-seq datasets from sucrose gradient-purified 50-80 bp subnucleosomes and from the fraction containing canonical nucleosomes allowed us to isolate the five clusters of putative 501 Mel cell enhancers shown in Extended Data Fig. 7.

#### List of CTCF-BS

We identified mESC genomic CTCF-BS based on the ChIP-seq signal of several CTCF datasets (GSM723015, GSM1828650, GSM288351 and GSM560352) at DNase-hypersensitive (DHS) regions, as described above for enhancers. All loci proximal (< 2 kb) to gene promoters and putative enhancers were removed. We used a subset (n = 9,850) of these CTCF-positive regions to generate the V-plots shown in Extended Data Fig. 8 and 9.

#### List of promoters

We rank-ordered gene promoters transcribed in mESCs according to the distance between the two divergent TSSs located on the plus and minus strands, using 5’ ends of Start-RNA reads (GSE43390), as described^66^. mESC Start-RNA reads datasets were first aligned to mm9 using Bowtie 1.3.0. We conserved only the reads mapping to a single position with a maximum of two mismatches. We derived the reference promoter list from Refseq annotation. We defined promoter windows as ± 1000 bases around the TSS position. We discarded windows with more than one TSS on the plus strand. We also removed the promoters associated with less than five overlapping Start-RNA reads. We calculated the coverage using a bin of one base. We selected the first six bases of the 5-prime reads as the signal to adjust the position of the TSS on the plus strand. We used a Python script to find the nearest local maximum upstream of the Refseq annotation, which we defined as the TSS position in this study. Next, we determined the location of the antisense Start-RNA signal on the minus strand in the region located upstream of the plus strand TSS. As mentioned above, we selected the first six bases of the 5-prime reads to define the position of the antisense TSS. We then designated the coordinate of the antisense TSS as the location having maximum Start-RNA expression on the minus strand, upstream of the sense TSS. We next sorted the promoters according to the distance between sense and antisense TSS coordinates. In this study, we focused our analysis on the 1,000 TSSs with the smallest distance between sense and antisense TSSs. The nucleosomal and subnucleosomal organization of the other groups of TSSs is similar and will be presented in detail in a separate publication.

### MNase ChIP-seq, MNase-seq, and ATAC-seq analysis

#### Quality controls, mapping and post-mapping process

The processes described below were done using Snakemake^67^, Conda and in-house Python scripts. We controlled the quality of sequencing data with FastQC (fastqc=0.11.9, default parameters)^68^ and fastqScreen (fastq-screen=0.14.0) using bowtie2 as default aligner^69^. Depending on their quality, sequencing files were trimmed with fastp (fastp=0.20.1) using the process described hereafter. Reads with a size below 25 bp were discarded to keep mapping specificity. Adapter sequences were auto-detected over ~1 million reads and removed. Reads with a base quality under 10 through a window size of 6 bases were discarded, and poly-G sequences were filtered out. Potential artefactual enrichments were assessed over 1/20^th^ of the raw reads and removed^70^. Trimmed paired-end reads were mapped with Bowtie 2 (bowtie2=2.4.2)^71^ over the Gencode’s 9^th^ mouse genome version for mESCs or the 38^th^ human genome version for 501 Mel cells. After mapping, reads were sorted by genomic coordinates using Samtools (samtools=1.11). Artefactual fragment duplication (optical and local) was assessed with Picard (picard=2.23.8; ‘Picard Toolkit.’ 2019. Broad Institute, GitHub Repository. http://broadinstitute.github.io/picard/; Broad Institute). Samtools was used to remove marked duplicates, unmapped, and reads involving a mapping quality below 30. The RPKM normalized genomic coverage was calculated with Deeptools for MNase data with a bin size of one base and centered on the fragment center for a fragment size of 10 to 500 bp. Chromosomes M, Y and X were not used for the normalization. The effective mouse genome size was set to 2,652,783,500 bp. (deeptools=3.5.0)^72^.

#### IGV visualizations and 2D histograms (V-plots)

Peaks and normalized coverage profiles were visualized as IGV (Integrative Genomics Viewer) screen-shots, heatmaps, and 2D graphs (V-plots). We converted BAM files containing paired-end mapped reads to BED files containing DNA fragments using Bedtools bamtobed (bedtools=2.29.2)^73^. We visualized these DNA fragments as a heatmap with seqMINER software and as a 2D graph (V-plots) using a homemade script. In the plot, the y-axis corresponds to DNA fragment size in bp and the x-axis to the relative position of DNA center fragments on the genome. Each dot corresponds to the coordinates of a DNA fragment midpoint, and color intensity reflects the number of fragments at that position. When comparing several samples, we chose the color intensity manually, and a ratio was applied to reduce the number of fragments to the same level in each sample.

### Analysis of the frequency at which subnucleosomal particles are organized in pairs at enhancers and determination of DNA linker length

IGV screen-shots of mESC enhancers from clusters 1 and 8 (n = 1,430) were analyzed individually and separated into three subgroups based on the enrichment pattern of histone H3, H4, H2A and H2B ChIP-seq peaks for 50-80 bp subnucleosomes: i) 496 enhancers displayed a single-module organization with two well-separated peaks, as shown in Fig. 4d and Extended Data Fig. 5; ii) 881 enhancers exhibited three or more discernible histone peaks, and were classified as multi-module enhancers (Extended Data Fig. 10); iii) 53 enhancers (3.7 % of total) presented a single histone peak that could be interpreted either as a rare subcategory of enhancers bearing a single subnucleosomal particle, or as single module enhancers having a pair of two closely located histone peaks fused into a single peak. For the three subgroups, the ChIP-seq peaks of histones H3, H4, H2A and H2B always overlapped in genomic position, confirming the systematic association of the four histones in these 50-80 bp subnucleosomes. We measured the distance D in bp between the centers of the two peaks of the subgroup of 496 single-module enhancers. Assuming that each of these two peaks marks the position of a hemisome protecting an average of 64 bp of DNA from MNase digestion, as proposed in Fig. 4f, we calculated the linker length between these two putative hemisomes as D minus 64 bp. We found that this DNA linker’s length ranges from 0 to 102 bp (median 29 bp) (Extended Data Fig. 5i).

### Detection of TF motifs at enhancers

We performed motifs analysis using the Meme suite (v5.3.0)^74^. Detection of OCT4 motifs in Fig. 4d, 5i-j, Extended Data Fig. 5, 10 and 11g was carried out with Fimo over selected lists of enhancers (clusters 1, 2, 3 and 8 from Extended Data Fig. 1), using two distinct p-values (0.001 and 0.01) to allow different levels of stringency in the motif sequence matching. The OCT4 motif matrix (<u>MA0142.1</u>) came from the Jaspar database^75^. We defined the OCT4, SOX2, NANOG and KLF4 motif enrichment profiles at mESC enhancers of clusters 1 and 2 using Centrimo (Extended Data Fig. 3b,c). We also used Centrimo to analyze the distribution of SOX10, MITF and TFAP2A^58^ motifs at putative enhancers in human 501 Mel cells (Extended Data Fig. 7)

### Detection of OCT4 full-size and half-site motifs in the DNA prepared from OCT4-immunopurified 50-80 bp subnucleosomes

OCT4 half-site motifs were taken from reference^11^. OCT4 full-size motifs and half-sites were detected with Fimo using two distinct p-values (0.001 and 0.01), as described above. The IGV screen-shots of enhancers from a representative cluster (cluster 8, n = 701) were analyzed individually to score the presence of both OCT4 full-size motifs and half-sites in the DNA of the 50-80 bp subnucleosomes bound by OCT4. These OCT4-enriched regions, which coincide with the histone peaks, are well defined on the IGV screen-shots (e.g. Fig. 4d, 5i-j, Extended Data Fig. 5 and 10). When we used the stringent p-value (p = 0.001), we found that 64% of the OCT4-enriched subnucleosomes did not contain an OCT4 full-size consensus motif nor a half-site. When we used the relaxed p-value (p = 0.01), allowing the detection of degenerated motifs and half-sites, we found that 18.5 % of the OCT4-bound subnucleosomes did not contain an OCT4 full-size motif nor a half-site. Note that a large majority of the OCT4 motifs detected with the relaxed p-value (p = 0.01) were not associated with a positive OCT4 ChIP-seq signal on histone-free DNA (e.g. Extended Data Fig. 5 and 10). It is thus unlikely that OCT4 binds these degenerated motifs onto 50-80 bp subnucleosomes *in vivo*. We concluded that a significant proportion of OCT4-bound subnucleosomes lack both OCT4 full-size motifs and half-sites in the DNA wrapped around the particle.

### Analysis of OCT4 occupancy at enhancers

For each enhancer of clusters 1 (n = 729), 2 (n = 600), 3 (n = 637) and 8 (n = 701), we first calculated the maximum rpkm value of the OCT4 ChIP-seq signal on the histone-free DNA fraction. Then, for each group of enhancers, we plotted the maximum rpkm values against the number of enhancers associated with each rpkm score. The curve revealed that most enhancers are associated with a low rpkm score, while a minority display a higher score. We identified a sharp bending of the curve (knee point or elbow) for each cluster. We choose the rpkm value at the knee point as the threshold defining a low and high OCT4 ChIP-seq signal. The rpkm threshold values were 1192, 1264, 1438, and 1294 for clusters 1, 2, 3, and 8, respectively. 64 (9%), 54 (9%), 76 (12%), and 71 (10%) enhancers of these clusters had a score above the threshold and were thus considered as having a high OCT4 ChIP-seq signal on histone-free DNA. Conversely, 665 (91%), 546 (91 %), 561 (88%), and 630 (90%) enhancers of clusters 1, 2, 3, and 8 had a low OCT4 ChIP-seq signal on histone-free DNA. Observations of IGV screen-shots confirmed that all enhancers having an rpkm above the threshold displayed well-identified OCT4 ChIP-seq peaks on histone-free DNA, whereas those below the threshold had either low or no detectable signal (e.g. Fig. 4d, 5i, Extended Data Fig. 5e). In contrast, the 50-80 bp subnucleosomes had a high OCT4 ChIP-seq signal at most enhancers. Inspection of IGV screen-shots of all enhancers of clusters 1, 2, 3 and 8 revealed that even those displaying the lowest rpkm scores had a well-defined OCT4 ChIP-seq enrichment signal. Thus, OCT4 occupancy is constitutively high on 50-80 bp subnucleosomes, in sharp contrast to its low occupancy of the OCT4 motifs present within histone-free DNA.

### TF motif detection in the DNA fragments purified from the top gradient fractions

We used MACS2 callpeak (macs2 2.2.7.1) to identify the genomic regions enriched in histone-free DNA fragments sequenced from the top gradient fractions (DNA purified from a pool of sucrose gradient 2-3-4). We set the minimal size for a peak to 40 bp. Then, motifs were detected with HOMER findMotifsGenome.pl for mouse genome (homer 4.11). For Extended Data Fig. 9g, we selected 32 TF motifs with a significant p-value (p < 10^−50^) in the samples prepared from mESCs expressing wild-type BRG1 (control samples). We noticed that the number of CTCF motifs detected was unaffected by the depletion of BRG1, in agreement with previous observations based on ATAC-seq experiments^76^. We thus choose to normalize the data according to the number of CTCF motifs detected in each sample. Using a homemade R script, the number of CTCF motifs found in control (wild-type BRG1) samples was divided by the number of CTCF motifs detected in BRG1-depleted samples. This coefficient was then applied to 31 other TF motifs to calculate the ratio between the number of motifs detected in chromatin samples prepared from BRG1-depleted and BRG1-expressing mESCs.

## Data availability

We deposited high-throughput sequencing data at the Gene Expression Omnibus with accession GSE210780, GSE210444 and GSE209914. Density graphs used in Fig. 4, 5 and Extended Data Fig. 5 and 10 have been deposited at the Zenodo Data repository: https://zenodo.org/record/7056534. The following public ATAC-seq datasets were obtained from GEO: mESCs depleted of OCT4 during 24h (GSM2341274, GSM2341275, GSM2341276, GSM2341284, GSM2341285, GSM2341286) and 15h (GSM5327548, GSM5327549, GSM5327538, GSM5327539).

## Acknowledgements

We thank A. Goldar, A. Leforestier, J.M. Victor, S. Khochbin, K. Yen, S. Mahony, I. Davidson for discussions, E. Nora and B. Bruneau for the E14Tg2a-Tir1 cell line, and C. Mann for reading the manuscript.

This work was supported by grants from the INCA (2017-1-PL BIO-02-CEA-1), Investissement d’Avenir ANR (Revive Labex ANR-10-LABX-73), ANR (ANR-17-CE12-0001-01), PRC/PICS07434 and Fondation ARC pour la recherche sur le cancer www.fondation-arc.org.

## Author’s contributions

M.C.N. designed, performed and analyzed experiments and bioinformatic analyses.

A.M.K. designed, performed and analyzed experiments.

E. D. achieved the majority of bioinformatic analyses.

W.A. performed and analyzed EM experiments.

C.D. and F.R. performed experiments.

J.A. performed and analyzed size exclusion chromatography experiments.

H.P. assisted with experiments.

O.A. performed bioinformatic analyses.

J.C.A. designed tools for bioinformatic analyses.

N.G. supervised EM experimentation and analysis

F.O. performed modelling of the split-nucleosome and contributed to data interpretation.

S.C. designed and analyzed experiments.

M.G. conceived the project, designed and analyzed experiments, and wrote the manuscript.

## Additional information (containing supplementary information line)

### Supplementary tables

Supplementary Table 1: antibodies used in this study

Supplementary Table 2: cell lines used in this study

Supplementary Table 3: plasmids used in this study

Supplementary Table 4: sequencing depth

