## Supplemental Tables 1-4 for "BRG1 generates subnucleosomes that expand OCT4 binding and function beyond DNA motifs at enhancers"

**ADDITIONAL INFORMATION (containing supplementary information line)**

**Supplementary tables:**

Supplementary Table 1: antibodies used in this study

Supplementary Table 2: cell lines used in this study

Supplementary Table 3: plasmids used in this study

Supplementary Table 4: sequencing depth

**Supplementary Table 1: antibodies used in this study**

| Target protein | Source | Reference | Company | Experiment |
|---|---|---|---|---|
| Histone H2A | Rabbit polyclonal | ab18255 | Abcam | MNase-ChIP-seq |
| Histone H2B | Rabbit polyclonal | ab1790 | Abcam | MNase-ChIP-seq<br>sequential ChIP-seq |
| Histone H3 | Rabbit polyclonal | ab1791 | Abcam | MNase-ChIP-seq<br>Western blot |
| Histone H4 | Rabbit polyclonal | ab7311 | Abcam | MNase-ChIP-seq |
| H3K27ac | Rabbit polyclonal | ab4729 | Abcam | MNase-ChIP-seq |
| Nanog | Rabbit polyclonal | RCAB-001P | Reprocell | MNase-ChIP-seq |
| Oct4 | Goat polyclonal | sc-8628 | Santa Cruz Biotechnology | MNase-ChIP-seq |
| Oct4 | Mouse monoclonal C-10 | sc-5279 | Santa Cruz Biotechnology | MNase-ChIP-seq<br>Western blot |
| Oct4 | Goat polyclonal | MBS420786 | myBioSource | sequential ChIP-seq<br>MNase-ChIP-seq |
| RNA Pol II | Mouse monoclonal F-12 | sc-55492 | Santa Cruz Biotechnology | MNase-ChIP-seq |
| Sox2 | Goat polyclonal | sc-17320 | Santa Cruz Biotechnology | MNase-ChIP-seq |
| TBP | Mouse monoclonal | ab51841 | Abcam | MNase-ChIP-seq |
| Brg1 | Rabbit monoclonal | ab110641 | Abcam | Western blot |
| Flag epitope | Mouse monoclonal M2 | F1804 | Sigma-Aldrich | MNase-ChIP-seq |
| no known specificity | Rabbit polyclonal | ab37415 | Abcam | sequential ChIP-seq, isotype control |

**Supplementary Table 2: cell lines used in this study**

| Name | Gene | Tag | Parental cell line | Cell type | Reference |
|---|---|---|---|---|---|
| Brg1-3FTH | *Smarca4* | $(Flag)_3$-TEV-HA | 46C | Mouse ES | 23 |
| Chd4-3FTH | *Chd4* | $(Flag)_3$-TEV-HA | 46C | Mouse ES | 23 |

| | | | | | |
|---|---|---|---|---|---|
| E14Tg2a-Tir1 | *OsTIR1* | - | E14Tg2a | Mouse ES | 44 |
| Smarca4-mAID | *Smarca4* | mAID | E14Tg2a-Tir1 | Mouse ES | This study |
| 501 MEL | _ | _ | _ | Human melanoma | 41 |

**Supplementary Table 3: plasmids used in this study**

| Name | Reference |
|---|---|
| pHYPER-shChd4 | 23 |
| pHYPER-Linker | 23 |
| pHYPER-shSmarca4-O5 | This study |
| pHYPER-shSmarca4-O7 | This study |
| pMK287 | Addgene #72825, reference 28 |
| pL452 | 23 |
| pMK287-Smarca4-mAID | This study |
| pL452-Smarca4-mAID | This study |
| pSpCas9(BB)-2A-Puro (PX459) | Addgene #62988, reference 43 |
| pSpCas9(BB)-2A-Puro-Smarca4-sgRNA1 | This study |
| pSpCas9(BB)-2A-Puro-Smarca4-sgRNA2 | This study |

**Supplementary table 4: Sequencing depth**

| Sample_Name | Total_Reads | Uniquely_Mapped_Reads | Cell_Type |
|---|---|---|---|
| Histone H3 MNase ChIP-seq total chromatin replicate 1 | 39405340 | 28943037 | Mouse ES |
| Histone H3 MNase ChIP-seq total chromatin replicate 2 | 60914784 | 44691604 | Mouse ES |
| Histone H3 MNase ChIP-seq total chromatin replicate 3 | 104873719 | 75863839 | Mouse ES |
| Histone H3 MNase ChIP-seq total chromatin 4X MNAse replicate 1 | 207078535 | 150738704 | Mouse ES |
| Histone H3 MNase ChIP-seq total chromatin 4X MNAse replicate 2 | 61781586 | 44585938 | Mouse ES |
| Histone H3 MNase ChIP-seq total chromatin 4X MNAse replicate 3 | 43479728 | 30315453 | Mouse ES |
| Histone H3 MNase ChIP-seq total chromatin Brg1-depleted auxin 20h replicate 1 | 31798277 | 22650134 | Mouse ES |
| Histone H3 MNase ChIP-seq total chromatin Brg1-depleted auxin 20h replicate 2 | 151773914 | 108380042 | Mouse ES |

| | | | |
|---|---|---|---|
| Histone H3 MNase ChIP-seq total chromatin Brg1-depleted auxin 20h replicate 3 | 88909566 | 63780557 | Mouse ES |
| Histone H3 MNase ChIP-seq total chromatin Brg1-depleted auxin 3h replicate 1 | 215743211 | 151092796 | Mouse ES |
| Histone H3 MNase ChIP-seq total chromatin Brg1-depleted auxin 3h replicate 2 | 185894596 | 130328826 | Mouse ES |
| Histone H3 MNase ChIP-seq total chromatin shRNA O7 Brg1 replicate 1 | 135876674 | 90899551 | Mouse ES |
| Histone H3 MNase ChIP-seq total chromatin shRNA O7 Brg1 replicate 2 | 119411342 | 79944667 | Mouse ES |
| Histone H3 MNase ChIP-seq total chromatin shRNA O5 Brg1 replicate 1 | 130868384 | 86086088 | Mouse ES |
| Histone H3 MNase ChIP-seq total chromatin shRNA O5 Brg1 replicate 2 | 39867879 | 26716226 | Mouse ES |
| Histone H3 MNase ChIP-seq total chromatin control shRNA replicate 1 | 34232952 | 23514406 | Mouse ES |
| Histone H3 MNase ChIP-seq total chromatin control shRNA replicate 2 | 42501184 | 28920463 | Mouse ES |
| Histone H3 MNase ChIP-seq total chromatin control shRNA replicate 3 | 69852580 | 45963957 | Mouse ES |
| Histone H3 MNase ChIP-seq total chromatin control shRNA replicate 4 | 44776510 | 29469046 | Mouse ES |
| Histone H3 MNase ChIP-seq total chromatin control shRNA replicate 5 | 54270556 | 36552039 | Mouse ES |
| Histone H3 MNase ChIP-seq total chromatin control shRNA replicate 6 | 44369423 | 30167964 | Mouse ES |
| Histone H3 MNase ChIP-seq total chromatin shRNA Chd4 replicate 1 | 65674590 | 43030059 | Mouse ES |
| Histone H3 MNase ChIP-seq total chromatin shRNA Chd4 replicate 2 | 59238286 | 39403166 | Mouse ES |
| Histone H2B MNase ChIP-seq total chromatin shRNA O7 Brg1 replicate 1 | 108056373 | 72482575 | Mouse ES |
| Histone H2B MNase ChIP-seq total chromatin shRNA O7 Brg1 replicate 2 | 112988489 | 72113344 | Mouse ES |
| Histone H2B MNase ChIP-seq total chromatin control shRNA replicate 1 | 137585584 | 88523196 | Mouse ES |
| Histone H2B MNase ChIP-seq total chromatin control shRNA replicate 2 | 121688126 | 80175008 | Mouse ES |
| Oct4 (sc-8628) MNase ChIP-seq total chromatin replicate 1 | 47707418 | 30322687 | Mouse ES |

| | | | |
|---|---|---|---|
| Oct4 (sc-8628) MNase ChIP-seq total chromatin replicate 2 | 59335509 | 37686806 | Mouse ES |
| Oct4 (sc-5279) MNase ChIP-seq total chromatin replicate 1 | 50067553 | 36557957 | Mouse ES |
| Oct4 (sc-5279) MNase ChIP-seq total chromatin replicate 2 | 45262372 | 32914621 | Mouse ES |
| Oct4 (sc-5279) MNase ChIP-seq total chromatin replicate 3 | 48004617 | 35248815 | Mouse ES |
| Sox2 MNase ChIP-seq total chromatin replicate 1 | 32627037 | 22232616 | Mouse ES |
| Sox2 MNase ChIP-seq total chromatin replicate 2 | 31389866 | 22504305 | Mouse ES |
| Nanog MNase ChIP-seq total chromatin replicate 1 | 30714584 | 20658681 | Mouse ES |
| Nanog MNase ChIP-seq total chromatin replicate 2 | 26928177 | 19395388 | Mouse ES |
| Nanog MNase ChIP-seq total chromatin replicate 3 | 28771749 | 21001937 | Mouse ES |
| TBP MNase ChIP-seq total chromatin replicate 1 | 49294798 | 34840023 | Mouse ES |
| TBP MNase ChIP-seq total chromatin replicate 2 | 66103656 | 24228508 | Mouse ES |
| RNA pol II MNase ChIP-seq total chromatin replicate 1 | 54324781 | 39977831 | Mouse ES |
| RNA pol II MNase ChIP-seq total chromatin replicate 2 | 36438128 | 25299559 | Mouse ES |
| Brg1 MNase ChIP-seq total chromatin replicate 1 | 115811642 | 77692626 | Mouse ES |
| Brg1 MNase ChIP-seq total chromatin replicate 2 | 25057246 | 15192424 | Mouse ES |
| H3K27ac MNase ChIP-seq total chromatin replicate 1 | 33263598 | 23911689 | Mouse ES |
| H3K27ac MNase ChIP-seq total chromatin replicate 2 | 21155550 | 9327529 | Mouse ES |
| H3K27ac MNase ChIP-seq total chromatin replicate 3 | 26421354 | 16001972 | Mouse ES |
| Sequential Oct4-H2B MNase ChIP-seq total chromatin replicate 1 | 139987435 | 97887367 | Mouse ES |
| Sequential Oct4-H2B MNase ChIP-seq total chromatin replicate 2 | 137731199 | 98766499 | Mouse ES |
| Sequential Control IgG-H2B MNase ChIP-seq total chromatin replicate 1 | 80698921 | 56390616 | Mouse ES |
| Sequential Control IgG-H2B MNase ChIP-seq total chromatin replicate 2 | 79867981 | 55784517 | Mouse ES |
| Histone H3 MNase ChIP-seq top of gradient (3-4) replicate 1 | 20473641 | 11040540 | Mouse ES |
| Histone H3 MNase ChIP-seq top of gradient (3-4) replicate 2 | 41170743 | 15263205 | Mouse ES |
| Histone H3 MNase ChIP-seq top of gradient (2-3-4) replicate 1 | 53669752 | 22469698 | Mouse ES |

| | | | |
|---|---|---|---|
| Histone H3 MNase ChIP-seq top of gradient (2-3-4) replicate 2 | 58774019 | 22497869 | Mouse ES |
| Histone H4 MNase ChIP-seq top of gradient (2-3-4) replicate 1 | 19352776 | 10225315 | Mouse ES |
| Histone H4 MNase ChIP-seq top of gradient (2-3-4) replicate 2 | 20614376 | 10173178 | Mouse ES |
| Histone H2B MNase ChIP-seq top of gradient (2-3-4) replicate 1 | 29435797 | 7712517 | Mouse ES |
| Histone H2B MNase ChIP-seq top of gradient (2-3-4) replicate 2 | 29458938 | 14052128 | Mouse ES |
| Histone H2A MNase ChIP-seq top of gradient (2-3-4) replicate 1 | 91847678 | 24503415 | Mouse ES |
| Histone H2A MNase ChIP-seq top of gradient (2-3-4) replicate 2 | 57971436 | 28515635 | Mouse ES |
| Histone H3 MNase ChIP-seq low sedimentation rate (5-6) replicate 1 | 293249523 | 189925120 | Mouse ES |
| Histone H3 MNase ChIP-seq low sedimentation rate (5-6) replicate 2 | 102514883 | 66497706 | Mouse ES |
| Histone H3 MNase ChIP-seq low sedimentation rate (5-6) replicate 3 | 38612558 | 27169665 | Mouse ES |
| Histone H3 MNase ChIP-seq low sedimentation rate (5-6) replicate 4 | 36081487 | 25887746 | Mouse ES |
| Histone H4 MNase ChIP-seq low sedimentation rate (5-6) replicate 1 | 52944646 | 37665907 | Mouse ES |
| Histone H4 MNase ChIP-seq low sedimentation rate (5-6) replicate 2 | 40526888 | 28765903 | Mouse ES |
| Histone H2B MNase ChIP-seq low sedimentation rate (5-6) replicate 1 | 40087420 | 31533332 | Mouse ES |
| Histone H2B MNase ChIP-seq low sedimentation rate (5-6) replicate 2 | 36132164 | 27808060 | Mouse ES |
| Histone H2A MNase ChIP-seq low sedimentation rate (5-6) replicate 1 | 47738304 | 19712186 | Mouse ES |
| Histone H2A MNase ChIP-seq low sedimentation rate (5-6) replicate 2 | 46769815 | 30031144 | Mouse ES |
| Histone H3 MNase ChIP-seq low sedimentation rate (5-6) - Brg1-depleted auxin 20h replicate 1 | 41330729 | 26810740 | Mouse ES |
| Histone H3 MNase ChIP-seq low sedimentation rate (5-6) - Brg1-depleted auxin 20h replicate 2 | 40869928 | 27019619 | Mouse ES |

| | | | |
|---|---|---|---|
| Histone H4 MNase ChIP-seq low sedimentation rate (5-6) - Brg1-depleted auxin 20h replicate 1 | 25719270 | 17162802 | Mouse ES |
| Histone H4 MNase ChIP-seq low sedimentation rate (5-6) - Brg1-depleted auxin 20h replicate 2 | 22724504 | 15100826 | Mouse ES |
| Histone H2B MNase ChIP-seq low sedimentation rate (5-6) - Brg1-depleted auxin 20h replicate 1 | 35490094 | 26948522 | Mouse ES |
| Histone H2B MNase ChIP-seq low sedimentation rate (5-6) - Brg1-depleted auxin 20h replicate 2 | 31999731 | 19556429 | Mouse ES |
| Histone H2A MNase ChIP-seq low sedimentation rate (5-6) - Brg1-depleted auxin 20h replicate 1 | 41922159 | 27850558 | Mouse ES |
| Histone H2A MNase ChIP-seq low sedimentation rate (5-6) - Brg1-depleted auxin 20h replicate 2 | 47392966 | 27265551 | Mouse ES |
| Histone H3 MNase ChIP-seq medium sedimentation rate (7-8) replicate 1 | 114190364 | 82909580 | Mouse ES |
| Histone H3 MNase ChIP-seq medium sedimentation rate (7-8) replicate 2 | 75002582 | 52035492 | Mouse ES |
| Histone H3 MNase ChIP-seq medium sedimentation rate (7-8-9) replicate 1 | 34805446 | 24044559 | Mouse ES |
| Histone H3 MNase ChIP-seq medium sedimentation rate (7-8-9) replicate 2 | 42053285 | 29950238 | Mouse ES |
| Histone H4 MNase ChIP-seq medium sedimentation rate (7-8-9) replicate 1 | 60490395 | 45691367 | Mouse ES |
| Histone H4 MNase ChIP-seq medium sedimentation rate (7-8-9) replicate 2 | 57404789 | 43374420 | Mouse ES |
| Histone H2A MNase ChIP-seq medium sedimentation rate (7-8-9) replicate 1 | 46891727 | 33652524 | Mouse ES |
| Histone H2A MNase ChIP-seq medium sedimentation rate (7-8-9) replicate 2 | 40975303 | 29674880 | Mouse ES |
| Histone H3 MNase ChIP-seq medium sedimentation rate (7-8-9) - Brg1-depleted auxin 20h replicate 1 | 43069748 | 28727096 | Mouse ES |
| Histone H3 MNase ChIP-seq medium sedimentation rate (7-8-9) - Brg1-depleted auxin 20h replicate 2 | 30354248 | 20344434 | Mouse ES |
| Histone H4 MNase ChIP-seq medium sedimentation rate (7-8-9) - Brg1-depleted auxin 20h replicate 1 | 61412390 | 43653168 | Mouse ES |

| | | | |
|---|---|---|---|
| Histone H4 MNase ChIP-seq medium sedimentation rate (7-8-9) - Brg1-depleted auxin 20h replicate 2 | 59980600 | 40742378 | Mouse ES |
| Histone H2A MNase ChIP-seq medium sedimentation rate (7-8-9) - Brg1-depleted auxin 20h replicate 1 | 48148096 | 32918764 | Mouse ES |
| Histone H2A MNase ChIP-seq medium sedimentation rate (7-8-9) - Brg1-depleted auxin 20h replicate 2 | 54422001 | 37321144 | Mouse ES |
| Histone H3 MNase ChIP-seq medium sedimentation rate (9-10) replicate 1 | 121992803 | 87067896 | Mouse ES |
| Histone H3 MNase ChIP-seq medium sedimentation rate (9-10) replicate 2 | 105877836 | 74113256 | Mouse ES |
| Histone H3 MNase ChIP-seq mononucleosomes (11) replicate 1 | 182771490 | 125965567 | Mouse ES |
| Histone H3 MNase ChIP-seq mononucleosomes (11) replicate 2 | 174543235 | 117406449 | Mouse ES |
| Histone H3 MNase ChIP-seq mononucleosomes (13) replicate 1 | 149358106 | 107675250 | Mouse ES |
| Histone H3 MNase ChIP-seq mononucleosomes (13) replicate 2 | 158934220 | 116584550 | Mouse ES |
| Oct4 (sc-5279) MNase ChIP-seq top of gradient (2-3-4) replicate 1 | 34609873 | 9948144 | Mouse ES |
| Oct4 (sc-5279) MNase ChIP-seq top of gradient (2-3-4) replicate 2 | 49817930 | 16033144 | Mouse ES |
| Oct4 (sc-5279) MNase ChIP-seq top of gradient (2-3-4) - Brg1-depleted auxin 20h replicate 1 | 36786141 | 15517232 | Mouse ES |
| Oct4 (sc-5279) MNase ChIP-seq top of gradient (2-3-4) - Brg1-depleted auxin 20h replicate 2 | 41246122 | 10929875 | Mouse ES |
| Oct4 (sc-5279) MNase ChIP-seq low sedimentation rate (5-6) replicate 1 | 41244905 | 28960438 | Mouse ES |
| Oct4 (sc-5279) MNase ChIP-seq low sedimentation rate (5-6) replicate 2 | 45435928 | 32366694 | Mouse ES |
| Oct4 (sc-5279) MNase ChIP-seq low sedimentation rate (5-6) - Brg1-depleted auxin 20h replicate 1 | 17921526 | 11896439 | Mouse ES |
| Oct4 (sc-5279) MNase ChIP-seq low sedimentation rate (5-6) - Brg1-depleted auxin 20h replicate 2 | 49705177 | 32221071 | Mouse ES |

| | | | |
|---|---|---|---|
| Oct4 (MBS420786) MNase ChIP-seq low sedimentation rate (5-6) replicate 1 | 57187594 | 41801325 | Mouse ES |
| Oct4 (MBS420786) MNase ChIP-seq low sedimentation rate (5-6) replicate 2 | 52195444 | 37436649 | Mouse ES |
| MNase-seq top of gradient (2-3-4) replicate 1 | 26003009 | 13283808 | Mouse ES |
| MNase-seq top of gradient (2-3-4) replicate 2 | 28738776 | 14029338 | Mouse ES |
| MNase-seq top of gradient (2-3-4) replicate 3 | 22599472 | 12471528 | Mouse ES |
| MNase-seq top of gradient (2-3-4) replicate 4 | 23155019 | 12098440 | Mouse ES |
| MNase-seq top of gradient (2-3-4) - Brg1-depleted auxin 20h replicate 1 | 21507231 | 10514555 | Mouse ES |
| MNase-seq top of gradient (2-3-4) - Brg1-depleted auxin 20h replicate 2 | 21099830 | 10371369 | Mouse ES |
| MNase-seq top of gradient (2-3-4) - Brg1-depleted auxin 20h replicate 3 | 23456104 | 10165690 | Mouse ES |
| MNase-seq top of gradient (2-3-4) - Brg1-depleted auxin 20h replicate 4 | 25998413 | 12657378 | Mouse ES |
| MNase-seq low sedimentation rate (5-6) replicate 1 | 24979900 | 15500698 | Mouse ES |
| MNase-seq low sedimentation rate (5-6) replicate 2 | 25385041 | 15778142 | Mouse ES |
| MNase-seq low sedimentation rate (5-6) replicate 3 | 25582206 | 16018924 | Mouse ES |
| MNase-seq low sedimentation rate (5-6) replicate 4 | 24255831 | 15269985 | Mouse ES |
| MNase-seq low sedimentation rate (5-6) - Brg1-depleted auxin 20h replicate 1 | 27360205 | 14856220 | Mouse ES |
| MNase-seq low sedimentation rate (5-6) - Brg1-depleted auxin 20h replicate 2 | 28305450 | 15260196 | Mouse ES |
| MNase-seq low sedimentation rate (5-6) - Brg1-depleted auxin 20h replicate 3 | 24603925 | 13974084 | Mouse ES |
| MNase-seq low sedimentation rate (5-6) - Brg1-depleted auxin 20h replicate 4 | 23757846 | 13642412 | Mouse ES |
| MNase-seq medium sedimentation rate (7-8-9) replicate 1 | 28798006 | 18729603 | Mouse ES |
| MNase-seq medium sedimentation rate (7-8-9) replicate 2 | 33555833 | 21747300 | Mouse ES |
| MNase-seq medium sedimentation rate (7-8-9) replicate 3 | 20381424 | 13336428 | Mouse ES |
| MNase-seq medium sedimentation rate (7-8-9) replicate 4 | 27531505 | 17681608 | Mouse ES |

| | | | |
|---|---|---|---|
| MNase-seq medium sedimentation rate (7-8-9) - Brg1-depleted auxin 20h replicate 1 | 21326975 | 11581715 | Mouse ES |
| MNase-seq medium sedimentation rate (7-8-9) - Brg1-depleted auxin 20h replicate 2 | 24582396 | 13317734 | Mouse ES |
| MNase-seq medium sedimentation rate (7-8-9) - Brg1-depleted auxin 20h replicate 3 | 25948329 | 14683588 | Mouse ES |
| MNase-seq medium sedimentation rate (7-8-9) - Brg1-depleted auxin 20h replicate 4 | 31301240 | 16610161 | Mouse ES |
| Histone H3 MNase ChIP-seq low sedimentation rate (4-5-6-7) replicate 1 | 70812228 | 46196630 | 501 Mel (human) |
| Histone H3 MNase ChIP-seq low sedimentation rate (4-5-6-7) replicate 2 | 71926960 | 44990018 | 501 Mel (human) |
| Histone H3 MNase ChIP-seq mononucleosomes (11) replicate 1 | 86713064 | 71542322 | 501 Mel (human) |
| Histone H3 MNase ChIP-seq mononucleosomes (11) replicate 2 | 105368488 | 87150338 | 501 Mel (human) |
